# BEAN and HABAS: Polyphyletic insertions in RNAP that point to deep-time evolutionary divergence of bacteria

**DOI:** 10.1101/2024.04.02.587612

**Authors:** Claudia Alvarez-Carreño, Angela T. Huynh, Anton S. Petrov, Christine Orengo, Loren Dean Williams

**Author notes:** Corresponding authors: Claudia Alvarez-Carreño, **Email:**, Loren Dean Williams **Email:**.

## Abstract

The β and β’ subunits of the RNA polymerase (RNAP) are large proteins with complex multi-domain architectures that include several insertional domains. Here, we analyze the multi-domain organizations of bacterial RNAP-β and RNAP-β’ using sequence, experimentally determined structures and AlphaFold structure predictions. We observe that bacterial lineage-specific domains in RNAP-β belong to a group of domains that we call BEAN (Broadly Embedded ANnex) and that in RNAP-β’, bacterial lineage-specific domains are HAmmerhead/BArrel-Sandwich Hybrid (HABAS) domains. The BEAN domain has a characteristic three-dimensional structure composed of two square bracket-like elements that are antiparallel relative to each other. The HABAS domain contains a four-stranded open β-sheet with a GD-box-like motif in one of the β-strands and the adjoining loop. The BEAN domain is identified not only in the bacterial RNAP-β’, but also in the archaeal version of universal ribosomal protein L10. The HABAS domain is observed as an insertional domain in several metabolic proteins. The phylogenetic distributions of bacterial lineage-specific insertional domains of β and β’ subunits of RNAP follow the Tree of Life. The presence of insertional domains can help establish a relative timeline of events in the evolution of a protein because insertion is inferred to post-date the base domain. We discuss mechanisms that might account for the discovery of homologous insertional domains in non-equivalent locations in bacteria and archaea.

## Introduction

Transcription is the Central Dogma process in which RNA polymerase (RNAP) transcribes DNA into RNA (Hurwitz, et al. 1961). mRNA is then translated into protein in the ribosome. RNAP contains five subunits called α1, α2, β, β’, and ϖ. The β and β’ subunits of RNAP, the focus of this work, are large proteins with complex multi-domain architectures. RNAP-β and RNAP-β’ both contain double-Ψ-β-barrel (DΨBB) domains, which combine to form the catalytic core of RNAP (Castillo, et al. 1999; Iyer, et al. 2003).

RNAP-β and RNAP-β’ have bacterial, archaeal, and eukaryotic orthologs and contain universal sequence motifs and domains (Sweetser, et al. 1987; Jokerst, et al. 1989; Lane and Darst 2010a, b). However, the domain architectures of RNAP-β and RNAP-β’ vary significantly between archaea and bacteria, and among bacteria. Archaea-specific domains of RNAP are conserved in eukaryotes (Figures S2-S3). Here, we use sequences and structures to begin to reconstruct an extraordinary succession of events during the deep evolutionary history of RNAP.

We observe that bacterial lineages acquired specific types of insertional domains at multiple locations of RNAP-β and RNAP-β’. Archaeal lineages acquired different insertional domains. The acquisition of these insertions occurred during the deep history of RNAP, billions of years ago. The results here allow us to establish a classification system for bacterial RNAP-β and RNAP-β’ subunits, based on type, location and chronology of domain insertion.

Some bacteria-specific domains of RNAP are observed only in certain bacterial lineages (Borukhov, et al. 1991; Severinov, et al. 1992; Iyer, et al. 2004; Lane and Darst 2010b, a; Huang, et al. 2015; Qayyum, et al. 2024). We use a naming scheme in which domains of RNAP-β and RNAP-β’ that occur in all archaea but not in bacteria are called here “a-specific” domains. Domains that occur in all bacteria but not in archaea are called “b-specific”. Domains that occur in some bacterial lineages but not others are called “b/lineage-specific”.

Proteins most commonly acquire new domains by terminal addition (Weiner, et al. 2006; Marsh and Teichmann 2010), generating tandem multidomain architectures. Yet, both RNAP-β and RNAP-β’ acquired domains by internal insertion. Insertional domains are less frequent than terminally-added domains (Manriquez-Sandoval and Fried 2022). In bacterial RNAPs, insertional domains have accreted on preexisting base domains and on each other. These accretion processes reveal dependencies that can be particularly useful in establishing timelines of evolutionary events; it is assumed that an insertional domain was acquired more recently than the domain that hosts it. Most of the a-, b- and b/lineage-specific domains of RNAP-β and RNAP- β’ are insertional. In multiple sequence alignments (MSAs), a-, b- and b/lineage-specific domains are observed as “blocks” (Vishwanath, et al. 2004). In MSAs of RNAP-β and RNAP-β’, universal domains are broken by these blocks (Figure S1).

We observe a broadly distributed b/lineage-specific insertional domain with idiosyncratic positions in RNAP-β. We call this domain BEAN (Broadly Embedded ANnex). We observe a b/lineage-specific insertional domain with idiosyncratic position in RNAP-β’. We call this domain HABAS (HAmmerhead/BArrel-Sandwich Hybrid). The BEAN domain is also identified in the archaeal version of universal ribosomal protein L10 (uL10). The HABAS domain is observed as an insertional domain in several metabolic proteins.

The locations and phylogenetic distributions of insertional domains in RNAP-β, RNAP-β’ and uL10 report on events that occurred in the deep evolutionary past. These insertional domains appear in distinct positions in the most deeply rooted bacterial lineages. We explore possible scenarios by which BEAN was idiosyncratically inserted in bacterial RNAP but not archaeal RNAP and was universally inserted in archaeal uL10 but not bacterial uL10. Here, we observe the results of processes that reshaped the multidomain architecture of bacterial and archaeal orthologs and tapered off after early evolution.

## Results

### Domain organizations of RNAP β and β’

The MSAs of RNAP-β and RNAP-β’ display block structures indicating universal as well as a-specific, b-specific and b/lineage-specific elements (Figures 1a and 2a). We analyzed the RNAP- β and RNAP-β’ domain organizations using orthologous sequences from a subsample of a reference set of evenly sampled bacterial genomes (Zhu, et al. 2019). The subsample used here includes representatives from all known major bacterial species and has been adapted from (Moody, et al. 2022). The blocks within the MSA were annotated using CATH (Sillitoe, et al. 2021) (Tables 1 and 2). The location and type of insertion were verified using experimentally determined structures (Berman, et al. 2000) and AlphaFold structure predictions (Jumper, et al. 2021; Varadi, et al. 2022). We observe small insertions (< 50 residues) that lack sequence similarity to each other or to domain entries in three classification databases: CATH (Sillitoe, et al. 2021), ECOD (Schaeffer, et al. 2017) and SCOPe (Chandonia, et al. 2017). These insertions are omitted from the analysis here.

**Table 1.**
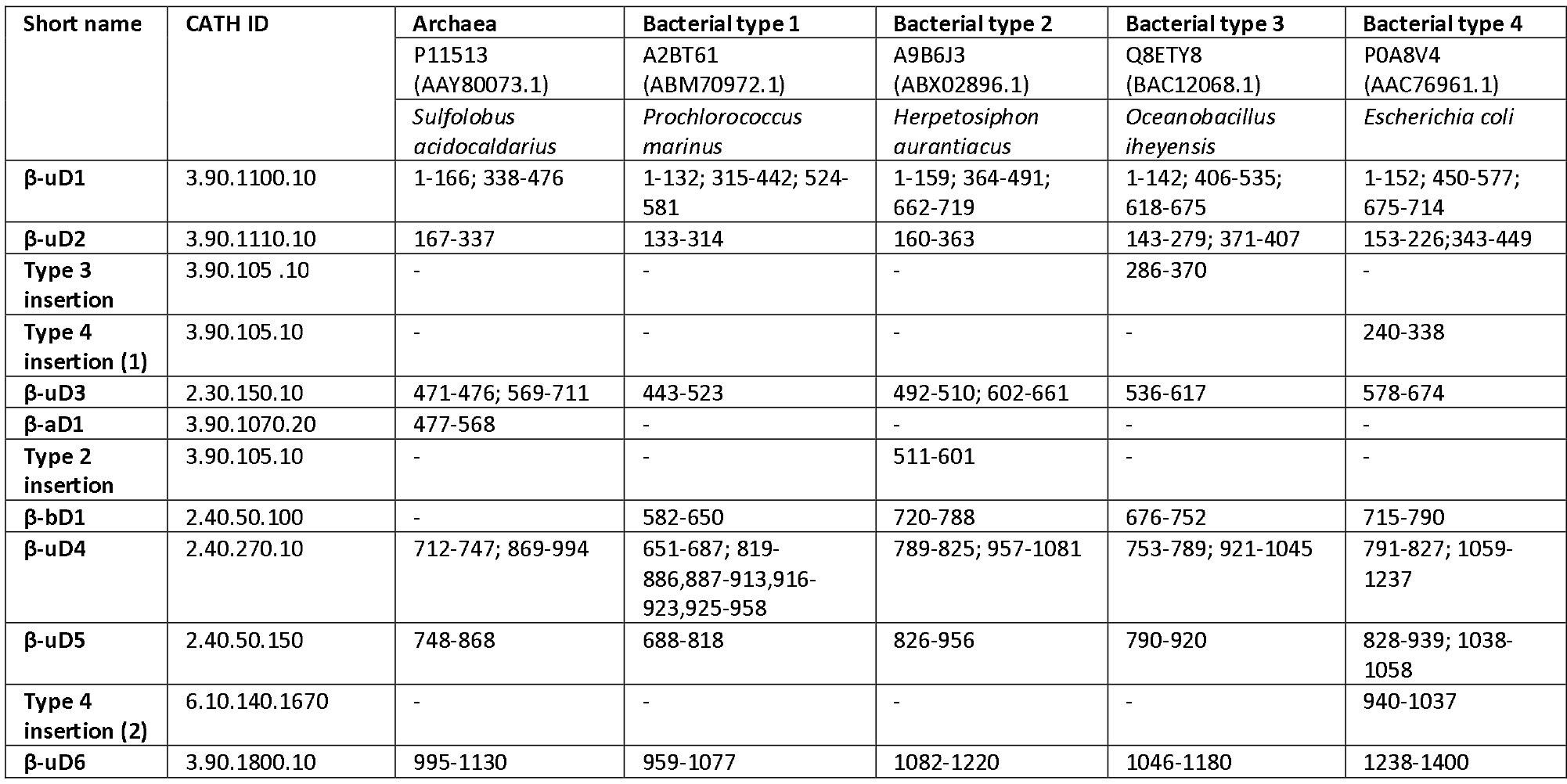
Multi-domain architecture of RNAP β subunit in representatives from Archaea and Bacteria

**Table 2.** Multi-domain architecture of RNAP β’ subunit in representatives from Archaea and Bacteria

| Short name | CATH | Archaea | Bacterial type 1 | Bacterial type 2 | Bacterial type 3 (N-ter) | Bacterial type 3 (C-ter) | Bacterial type 4 |
| --- | --- | --- | --- | --- | --- | --- | --- |
|  |  | P11512<br>(AAY80072.1) | Q0AUH3<br>(ABI69631.1) | Q3Z8V3<br>(AAW40096.1) | A2BT59<br>(ABM70970.1) | A2BT60<br>(ABM70971.1) | A7IKQ1<br>(ABS68594.1) |
|  |  | *Sulfolobus acidocaldarius* | *Syntrophomonas wolfei* subsp. *wolfei* | *Dehalococcoides mccartyi* | *Prochlorococcus marinus* | *Prochlorococcus marinus* | *Xanthobacter autotrophicus* |
| $\beta'$-uD1 | 4.10.960.120 | 1-148; 176-317 | 10-64; 93-138; 178-343 | 1-56; 84-130; 237-402 | 1-70; 98-144; 192-357 | - | 1-67; 95-141; 182-348 |
| $\beta'$-bD1 | 3.90.820.30 | - | 65-91 | 57-83 | 71-97 | - | 68-94 |
| a- insertion (1) | - | 149-175 | - | - | - | - | - |
| $\beta'$-bD2 | 3.30.60.280 | - | 139-177 | 131-140; 211-236 | 145-191 | - | 142-181 |
| Type 2 insertion | - | - | - | 141-210 | - | - | - |
| $\beta'$-uD2 | 2.40.40.20 | 318-346; 414-494 | 344-371; 414-495 | 403-430; 475-554 | 358-385; 428-509 | - | 349-376; 419-500 |
| $\beta'$-bD3 | 1.10.40.90 | - | 372-413 | 431-474 | 386-427 | - | 477-418 |
| $\beta'$-aD1 | 3.30.1490.180 | 347-413 | - | - | - | - | - |
| $\beta'$-uD3 | 1.10.274.100 | 495-560; 599-644 | 496-512; 579-640 | 555-571; 618-679 | 510-526; 618-635 | - | 501-517; 572-637 |
| $\beta'$-bD4 | - | - | 513-578 | 572-617 | 527-617 | 1-84 | 518-571 |
| $\beta'$-aD2 | 2.60.40.2940 | 561-598 | - | - | - | - | - |
| $\beta'$-uD4 | 1.10.132.30 | 645-832 | 641-798 | 680-837 | - | 85-244 | 638-806 |
|  |  | 1054-1093 |  |  |  |  |  |
| a- insertion (2) | - | 756-779 | - | - | - | - | - |
| $\beta'$-bD5 | 3.90.105.10 | - | 799-879 | 838-917 | - | 245-318 | 807-888 |
| $\beta'$-uD5 | 1.10.1790.20 | - | 880-949; 1013-1059; 1104-1113 | 918-987; 1118-1164; 1209-1282 | - | 319-369; 1009-1027; 1092-1154; 1199-1208 | 889-939; 1124-1143; 1208-1254; 1299- |
|  |  |  |  |  |  |  | 1308 |
| **β'-bD6** | 2.40.50.100 | - | 950-1012 | 988-1117 | - | 1028-1091 | 1144-1207 |
| **Type 3**<br>**insertion** | 2.40.50.100 | - | - | - | - | 370-1008* | - |
| **Type 4**<br>**insertion** | 1.10.132.30 | - | - | - | - | - | 940-1123* |
| **β'-bD7** | 3.30.60.280 | - | 1060-1103 | 1165-1208 | - | 1155-1198 | 1255-1298 |
| **β'-uD6** | 1.10.150.390 | - | 1114-1181 | 1219-1283 |  | 1209-1252 | 1309-1375 |

**Figure 1.**
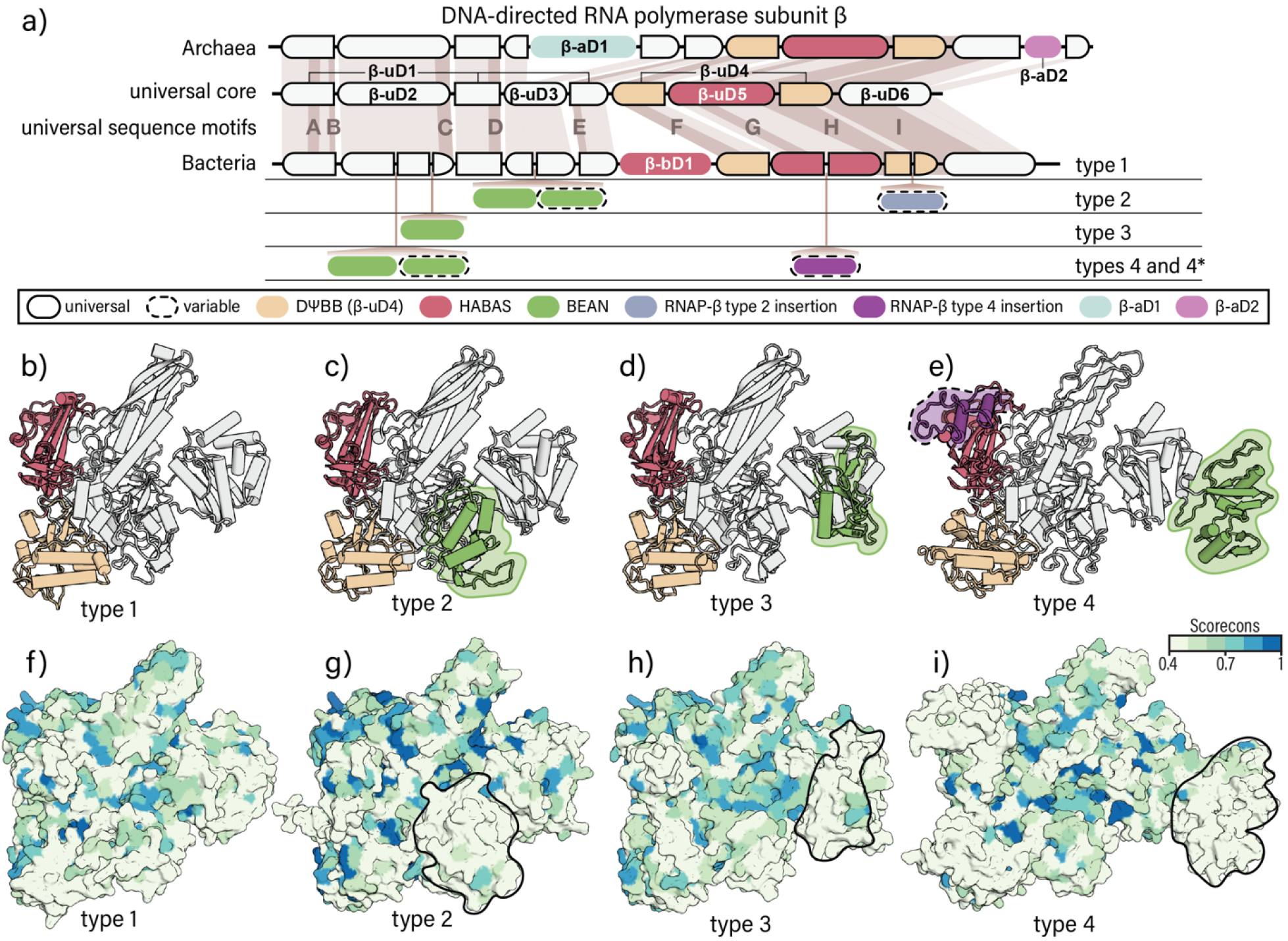
Multi-domain organizations of RNAP-β. (a) Multi-domain organization diagrams of RNAP-β in archaea and bacteria. First row: domains of archaeal RNAP-β. Second row: universally shared domains between archaeal and bacterial orthologs and universal sequence motifs described in (Sweetser, et al. 1987). Third row: domains of bacterial type 1 RNAP-β. Fourth row: location of bacterial type 2 insertions. Fifth row: location of bacterial type 3 insertions. Sixth row: location of bacterial type 4 insertions (b to e) Three-dimensional structure of the bacterial RNAP-β. Domains are coloured as in (a). (b) type 1 (AlphaFold DB: AF-A2BT61-F1), (c) type 2 (AlphaFold DB: AF-A9B6J3-F1), (d) type 3 (AlphaFold DB: AF-Q8ETY8-F1), and (e) type 4 (PDB: 4IGC, chain C). (f to i) Bacterial RNAP-β coloured by residue conservation. The conservation score ranges from 0 (not conserved) to 1 (highly conserved). Valdar01 score was calculated on the multiple sequence alignment of sequence representatives for each type using Scorecons. For clarity, N- and C-terminal residues that extend beyond the shared core of bacteria were masked.

**Figure 2.**
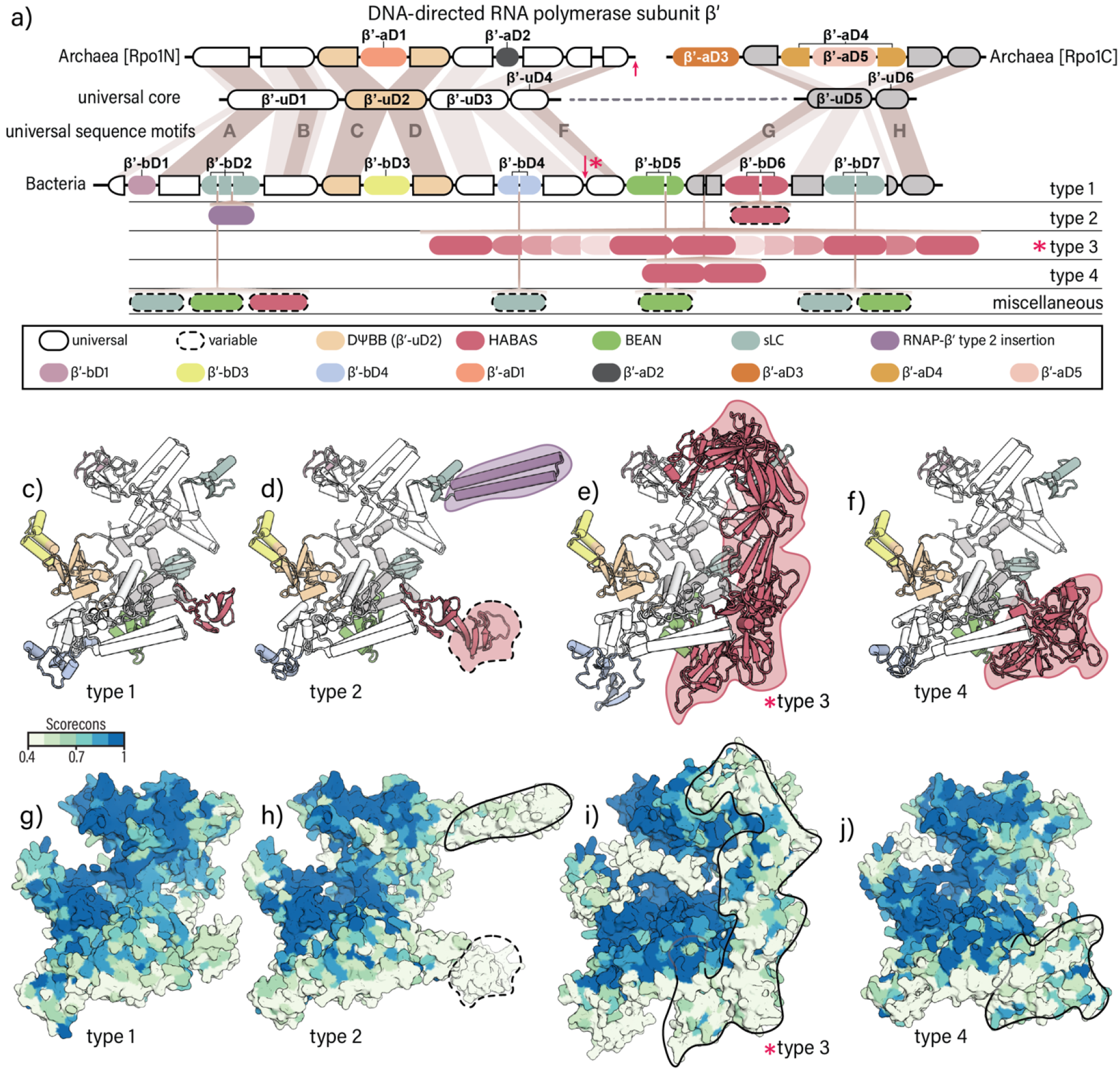
Multi-domain organization of RNAP-β’. (a) Multi-domain organization diagrams of RNAP-β’ in archaea and bacteria. First row: domains of archaeal orthologs. Second row: universally shared domains between archaeal and bacterial orthologs and universal sequence motifs described in (Jokerst, et al. 1989). Third row: domain organization of the bacterial type 1 RNAP-β’. Fourth row: location of bacterial type 2 insertions. Fifth row: location of bacterial type 3 insertions. Sixth row: location of bacterial type 4 insertions. Red arrows: sites of split of RNAP-β’ into sub-subunits. (b to e) Three-dimensional structures of bacterial RNAP-β’ colored by domains as in (a). (b) type 1 (AlphaFold DB: AF-Q0AUH3-F1), (c) type 2 (AlphaFold DB: AF-Q3Z8V3-F1), (d) type 3, N-terminal fragment (AlphaFold DB: AF-A2BT59-F1) and C-terminal fragment (AlphaFold DB: AF-A2BT60-F1), and (e) type 4 (AlphaFold DB: AF-A7IKQ1-F1). *Site of truncation of bacterial type 3 RNAP-β. (f to i) Bacterial RNAP-β’ coloured by residue conservation. The conservation score ranges from 0 (not conserved) to 1 (highly conserved). Valdar01 score was calculated on the multiple sequence alignment of sequence representatives for each type using Scorecons. For clarity, N- and C-terminal residues that extend beyond the shared core of bacteria were masked.

Our inquiry here is facilitated by our naming system for RNAP domains. In this scheme, the subunit is indicated by β or β’, followed by a hyphen and the letter “u” to indicate universal conservation, or “a” to indicate a-specific, or “b” to indicate b-specific. The domains (D) are numbered in order of appearance in the sequence. For example, the N-terminal RNAP β-subunit domain, which is universal, is called β-uD1.

### RNAP-β multi-domain architectures.

RNAP-β has six universal domains that are conserved in archaeal and bacterial orthologs. These domains are β-uD1, β-uD2,… β-uD6 (Figure 1). Universal insertional domains were identified at three different locations of RNAP-β: β-uD2 is insertional in β-uD1; β-uD3 is insertional in β-uD1; and β-uD5 (a HABAS domain) is insertional in β-uD4 (the DΨBB domain).

Archaea and bacteria RNAP-βs each contain additional domains (Figure 1a). In archaea, β-aD1 is inserted within β-uD3 and β-aD2 is inserted within β-uD6. In bacteria, β-bD1 is inserted between β-uD2 and β-uD4.

RNAP-β contains b/lineage-specific domains at multiple positions (Figure 1a). These domains are inserted at five distinct sites of RNAP-β: in the i) central region of β-uD2; ii) C-terminal half of β-uD2; iii) N-terminal half of β-uD3; iv) C-terminal half of β-uD5; and v) C-terminal half of β-uD4. The b- and b/lineage-specific domains in RNAP-β are less conserved in sequence than universal domains (Figure 1f to 1i). The b/lineage-specific insertions in β-uD2 and in β-uD3 share sequence and structure similarity (Figures 3c and 3d), suggesting common ancestry. We call these domains BEAN (Broadly Embedded ANnex).

**Figure 3.**
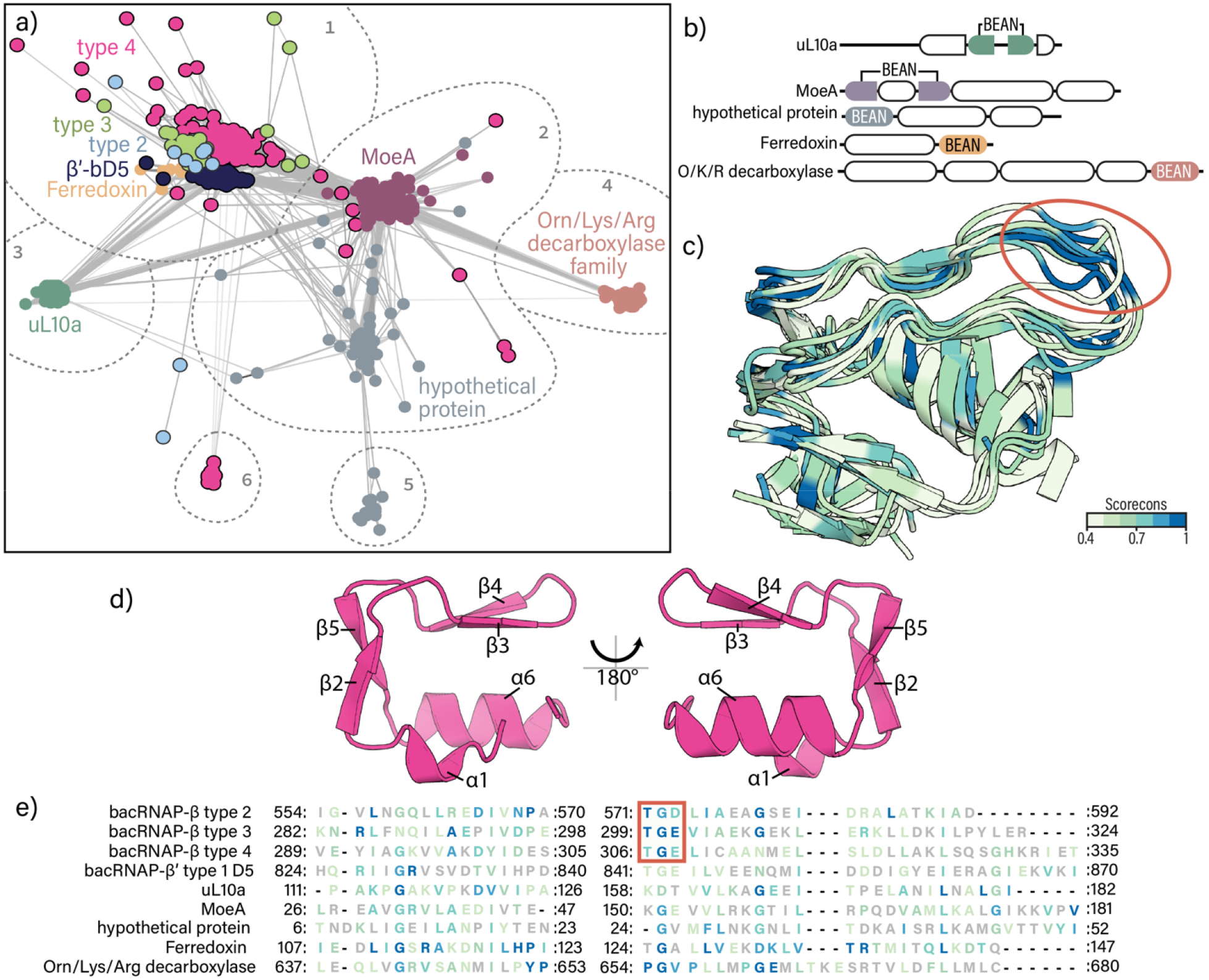
The BEAN domain. (a) BEAN domain sequences clustered by similarity at a P-value threshold of 1×10^-11^. Dotted lines encircle sequence clusters that remain at a P-value of 1×10^-15^. (b) Multi-domain organization of representative proteins containing BEAN domains. (c) Structure superimposition of BEAN domains from representative predicted structures. Structures are colored by conservation. (d) Structure of the BEAN domain (AlphaFold DB: AF-P0A8V4-F1). (e) Structure-derived multiple sequence alignment of RNAP-β type 2 (AlphaFold DB: AF-), type 3 (AlphaFold DB: AF-Q8ETY8-F1) and type 4 (AlphaFold DB: AF-P0A8V4-F1); RNAP-β’ (AlphaFold DB: AF-Q0AUH3-F1); uL10a (AlphaFold DB: AF-Q8TZJ8-F1); MoeA (AlphaFold DB: AF-O59354-F1); a hypothetical protein (AlphaFold DB: AF-B8J6M3-F1); Ferredoxin (AlphaFold DB: AF-Q4C556-F1); and ornithine/lysine/arginine decarboxylase (AlphaFold DB: AF-P52095-F1). Above 0.5, residues are colored by conservation score. Below 0.4 residues are shown in gray.

Sites of BEAN insertion within RNAP-β define b/lineage-specific RNAP-β architecture (Figure 1a). Differential b/lineage insertion in RNAP-β allows us to distinguish four types of RNAP-β. Type 1 RNAP-β lacks b/lineage-specific insertions (Figure 1a and 1b). Type 2 RNAP-β has a BEAN domain inserted within β-uD3 (Figure 1a and 1c). Type 3 RNAP-β has a BEAN domain inserted within β-uD2 (Figure 1a and 1d). Type 4 RNAP-β has a BEAN domain inserted within β-uD2 (Figure 1a and 1e). Certain RNAP-β proteins display a type 4 multidomain architecture with additional insertions; these are referred to as type 4*.

#### The BEAN domain

The BEAN domain has a characteristic three-dimensional structure composed of two square bracket-like elements that are anti-parallel relative to each other (Figure 3d). Each bracket-like element is formed by an α-helix and two β-strands. The orientation between consecutive secondary elements is 90° within each bracket. The first bracket is formed by α1⊥β2⊥β3 and the second bracket by β3⊥β4⊥α5. The BEAN domain maps to CATH superfamily 3.90.105.10.

We have identified BEAN domains in bacterial and archaeal proteins other than RNAP-β (Figure 3a to 3d). Using sequence and structure similarities, we find the BEAN domain in bacterial RNAP-β’; the archaeal version, but not the bacterial version of ribosomal protein uL10; molybdenum cofactor biosynthesis protein MoeA; ornithine/lysine/arginine decarboxylases; a putative ferredoxin; and in one uncharacterized protein.

A cluster analysis of sequences based on BLASTP P-values shows that b/lineage-specific BEAN domains in RNAP-β are more similar to each other than to BEAN domains in other proteins (Figure 3a). Additionally, BEAN domains in RNAP-β show a conserved TGD/E (threonine, glycine, aspartic/glutamic acid) sequence motif (Figure 3d) that is absent from other BEAN domains. Thus, BEAN domains in RNAP-β appear to share more recent ancestry with each other than with BEAN domains of other proteins.

### RNAP-β’ multi-domain architectures.

Bacterial RNAP-β’ domains map to two distinct proteins in archaea (Rpo1N and Rpo1C). Rpo1N and Rpo1C assemble to form a complete RNAP-β’ in archaea and eukaryotes, which is also called Rpo1 (Korkhin, et al. 2009). The naming system for domains in the archaeal RNAP-β’ follows a continuous order from Rpo1N to Rpo1C. Domains β’-uD1 to β’-uD4 are common to bacterial RNAP-β’ and archaeal Rpo1N; and domains β’-uD5 and β’-uD6 are common to bacterial RNAP-β’ and archaeal Rpo1C (Figure 2a). The DΨBB domain of RNAP-β’ is β’-uD2.

We identified a-specific insertional domains within β’-uD2 and β’-uD3 (Figure 2) and small a-specific insertions in Rpo1N, within β’-uD1 and β’-uD4. We also identified three a-specific domains in Rpo1C: β’-aD3, β’-aD4 and β’-aD5. β’-aD3 is an N-terminal addition to Rpo1C; β’-aD4 is inserted into β’-uD5, and β’-aD5 is inserted into β’-aD4.

Bacterial RNAP-β’ is composed of six universal RNAP-β’ domains and seven b-specific domains (type 1, Figure 2). Domains β’-bD1, β’-bD2, β’-bD3, β’-bD4, β’-bD6 and β’-bD7 are insertional. β’-bD5 is a BEAN domain and β’-bD6 is a HABAS domain. β’-bD3 is inserted into the DΨBB domain of the β’ subunit and shows no sequence or structure similarity to the archaeal DΨBB domain insertion (β’-aD1). β’-bD2 and β’-bD7 are homologous (CATH superfamily 3.30.60.280). Here, we call these domains sLC (small Left Claw).

The b/lineage-specific insertions in RNAP-β’ are observed in seven distinct locations (Figure 2a): i) in the first half of β’-bD2; ii) in the middle of β’-bD2; iii) in the middle of β’-bD4; iv) in the second half of β’-bD5; v) near the N-terminus of β’-uD5; vi) in the second half of β’-bD6; and vii) in the middle of β’-bD7. Based on the presence and location of b/lineage-specific insertions, we have defined four types of bacterial RNAP-β’ subunits (Figure 2a). Type 1 RNAP-β’ in bacteria has no b/lineage-specific insertions. Type 2, 3 and 4 bacterial RNAP-β’ have b/lineage-specific insertional domains. Most of these b/lineage-specific insertions are HABAS domains, which share sequence and structural similarity (Figures 2d to 2f).

The location and number of b/lineage-specific insertions are used here to define the ‘type’ of RNAP-β’. Bacterial type 1 RNAP-β’ contains a single BEAN domain (β’-bD5) and single HABAS domain (β’-bD6). Bacterial type 2 RNAP-β’ contains a BEAN insertion within β’-bD2 and sometimes a HABAS domain within β’-bD6. Bacterial type 3 and 4 RNAP-β’ have HABAS domain insertions within RNAP β’-uD5. These insertions differ in number but not in location. For example, bacterial type 4 RNAP-β’ contains one HABAS domain whereas bacterial type 3 RNAP-β’ contains nine HABAS domains. It appears that type 3 RNAP-β’ is an elaboration of type 4 RNAP-β’. HABAS domains in type 3 RNAP-β’ are inserted into other HABAS domains (Chlenov, et al. 2005; Qayyum, et al. 2024). These recursively inserted domains in some cases exchange secondary structural elements (are domain swapped), forming complex interdigitated structures (Figure 2e). Beyond the well-populated types described here, eleven additional RNAP-β’ variants are observed, with various combinations of HABAS, BEAN and sLC domains inserted within β’-bD2, β’-bD4, β’-bD5 and β’-bD7. This collection includes RNAP-β’ architectures with one or a few (three or less) representatives in our dataset of species (Table S4). All of these species are in deeply rooted lineages such as Firmicutes and the DST group (Deinococcus-Thermus, Synergistetes, Thermotogae and related bacteria).

Note that bacterial type 3 RNAP-β’ is composed of two polypeptide chains. The N-terminal sub-subunit (RNAP-β’_BacN_) ends with β’-uD3; and the C-terminal sub-subunit (RNAP-β’_BacC_) starts at β’uD4 (Figure 2a). RNAP-β’_BacN_ and RNAP-β’_BacC_ assemble to form a complete RNAP-β’ (Qayyum, et al. 2024).

#### The HABAS domain

Many b/lineage-specific insertions in RNAP subunit β’ are HABAS domains (CATH superfamily 2.40.50.100). The HABAS domain is also observed in RNAP-β in archaea and bacteria as β-uD5, and as a b-specific domain (β-bD1). Sequence similarity searches reveal additional proteins containing HABAS domains. HABAS inserted proteins are involved in metabolism (phosphatidylserine decarboxylase proenzyme, biotin carboxyl carrier protein, ion-translocating oxidoreductase complex subunit C, Cytochrome f), transport (major facilitator superfamily transporter protein, Resistance-Nodulation-Division family transporter protein) and genetic information processing (Transcription termination/antitermination protein NusG) (Figure 4).

**Figure 4.**
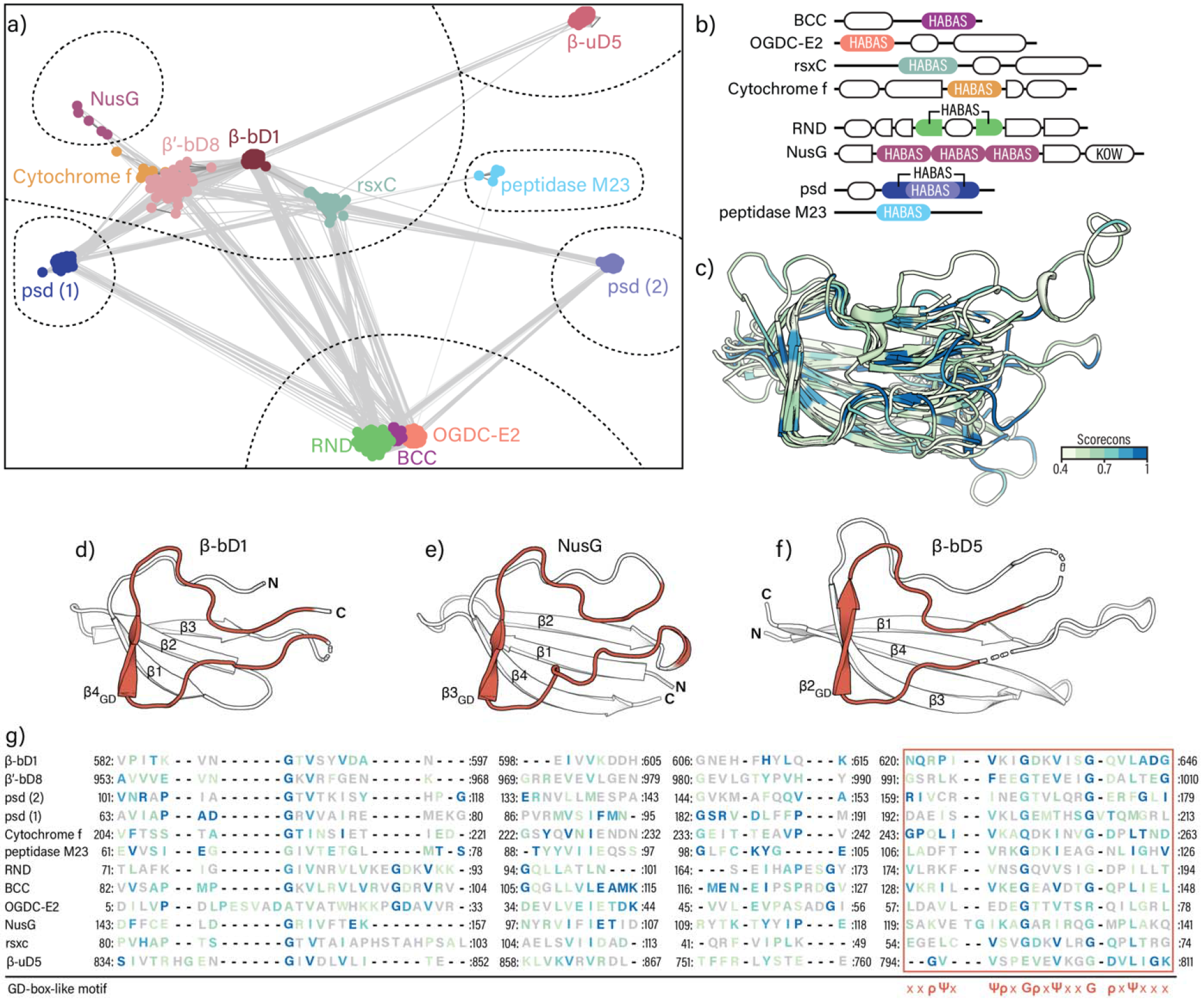
The HABAS domain. (a) HABAS domains clustered by sequences similarity at a P-value threshold of 5×10^-10^. Dotted lines encircle sequence clusters that remain at a P-value of 1×10^-12^. (b) Multi-domain organization of representative proteins containing HABAS domains. (c) Structure superimposition of HABAS domains from representative predicted structures. (d-e) Structure representation of HABAS domains. GD-box-like motif highlighted in dark orange. (d) HABAS domain β4_GD_ topology (AF-A2BPU4-F1). Residues 616-625 are masked. (e) HABAS domain β3_GD_ topology (AF-P77611-F1). (f) HABAS domain β2_GD_ topology (AF-C5CGE4-F1). Residues 711-725 and 747-775 are masked. (g) Structure-derived multiple sequence alignment of β-bD1 (A2BT61), β’-D8 (Q0AUH3), psd (A0L627), cytochrome f (A2BPU4), peptidase M23 (A0A0E3QUJ6), RND (A0A1T5M7J5), BCC (O59021), OGDC-E2 (P0AFG6), NusG (C5CGE4), rsxC (P77611) and β-uD5 (P11512). Above 0.5, residues are colored by conservation score. Below 0.4 residues are shown in gray.

The HABAS domain is a four-stranded open β-sheet with a conserved sequence motif in one of the β-strands and the adjoining loop. The motif contains glycine (G), aliphatic (Ψ), and polar (ρ) amino acids as follows: ΨxΨρxGρxΨxxGρxΨxx. We call this motif the GD-box-like motif because it is similar but not identical to the GD-box sequence motif ΨxΨxxGρxΨxΨ (Alva, et al. 2009). We found three distinct topologies of secondary structural elements in HABAS domains. These topologies are related by circular permutation. We distinguish and name these topological variants by the locations of their GD-box-like motifs. The most frequently observed topology has a GD-box-like motif in strand β4 (β4_GD_). β-bD1 has a β4_GD_ topology (Figure 4d). β-uD5 has a β2_GD_ topology (Figure 4e), and a HABAS domain of NusG has a β3_GD_ topology (Figure 4f).

### Phylogenetic distribution of RNAP-β and RNAP-β’ types

The phylogenetic distribution of RNAP-β and RNAP-β’ types as we define them here follows the deeply rooted divergence of Gracilicutes and Terrabacteria (Coleman, et al. 2021; Witwinowski, et al. 2022). We identified type 4 RNAP-β and type 4 RNAP-β’ in most Gracilicutes; by contrast, we identified all RNAP-β and RNAP-β’ types in Terrabacteria (Figure 5). The β and β’ subunits are fused into one polypeptide chain in Candidatus *Adlerbacteria* and *Wolinella succinogenes* (Table S1).

**Figure 5.**
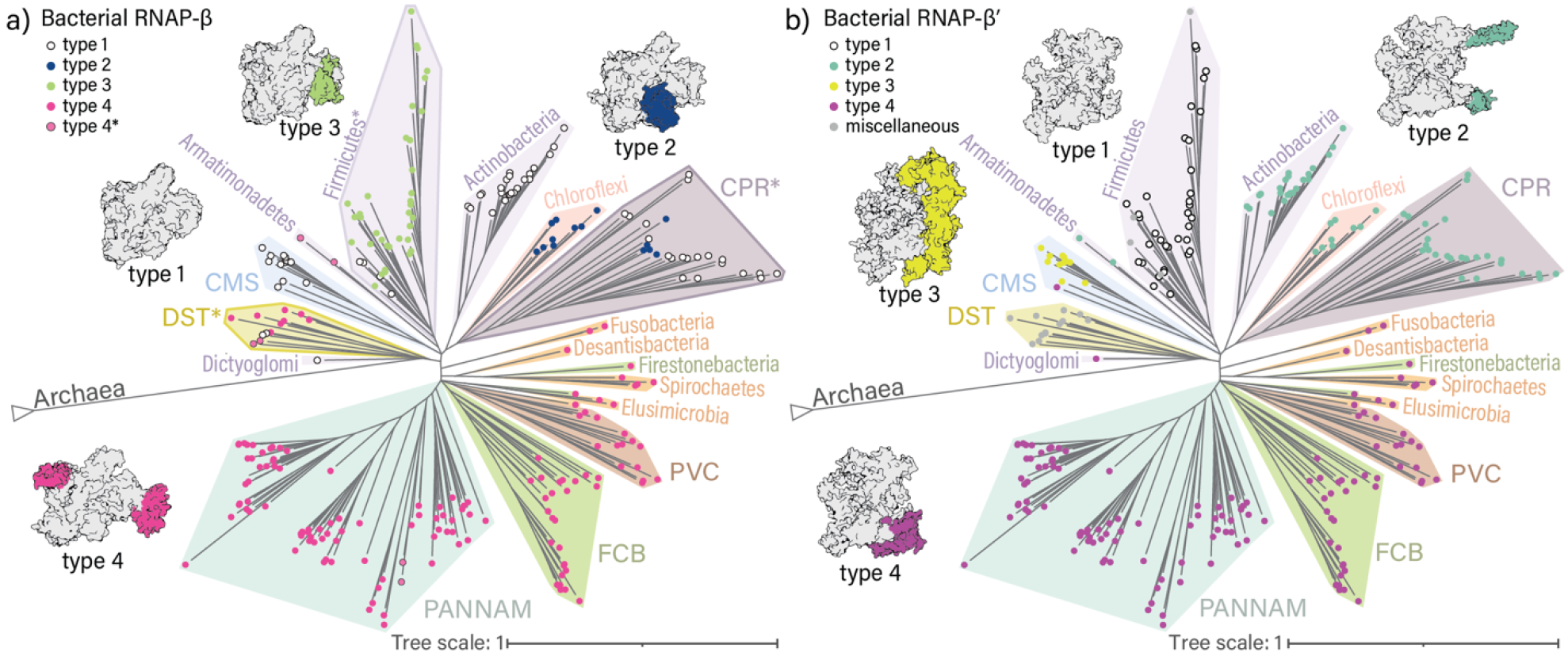
Phylogenetic distribution of the different multi-domain organizations of RNAP-β and RNAP-β’ in bacteria. Domain organization types are shown in different colors. The tree of Bacteria was adapted from (Moody, et al., 2022). (a) Distribution of RNAP-β types in bacteria. Phylogenetic groups with a scattered distribution of type 1 RNAP-β are indicated by a darker outline. (b) Distribution of RNAP-β types in bacteria. CMS: Cyanobacteria, Margulisbacteria, Melainabacteria; CPR: Candidatus Phyla Radiation; DST: Deinococcus-Thermus, Synergistes, Thermotogae, Bipolaricaulota, Caldiserica, Coprothermobacterota; FCB: Fibrobacteres, Chlorobi, Bacteroides, Gemmatimonadetes, Candidatus Cloacimonetes, division KSB1, Eisenbacteria, Candidatus Fermentibacteria, Firestonebacteria, Candidatus Glassbacteria, Ignavibacteria, Kryptonia, Marinimicrobia, Raymondbacteria, Stahlbacteria, Zixibacteria; PANNAM: Bdellovibrio, Dependentia, Proteobacteria, Aquificae, Myxococcota, Nitrospinae, Nitrospirae, Acidobacteria, Chrysiogenetes, Deferribacteres, Schekmanbacteria and Thermodesulfobacteria; PVC: Planctomycetes, Verrucomicrobia, Chlamydiae, Kiritimatiellaeota, Lentisphaerae, Candidatus Desantisbacteria, Candidatus Omnitrophica.

A given bacterial lineage tends to have a single type of RNAP-β. Bacteria in the CMS group (Cyanobacteria and related bacteria) contain type 1 RNAP-β; Armatimonadetes contain type 4* RNAP-β; Actinobacteria contain type 1 RNAP-β; and Chloroflexi contain type 2 RNAP-β. However, type 1 is observed mixing among other types of RNAP-β in some bacterial lineages. Most bacteria in the DST group contain type 4 RNAP-β, but some contain type 1. Firmicutes generally contain type 3 RNAP-β, but some contain type 1. Bacteria in the CPR group (Candidate Phyla Radiation) contain either type 2 or type 1 RNAP-β.

Type 1 RNAP-β lacks b/lineage-specific BEAN insertions. Its scattered phylogenetic distribution could correspond to HGT or reduction from more elaborate types. To test whether the scattered distribution of type 1 RNAP can be attributed to HGT, we calculated a maximum likelihood gene tree of RNAP-β using sites that are conserved in all bacteria (Figure 6). We compared the gene tree of RNAP-β to a consensus tree of bacteria calculated previously using 27 vertically inherited genes (Moody, et al. 2022). Our phylogenetic analysis shows that the tree of RNAP-β follows the the tree of bacteria (Figures 6 and S4) and suggests vertical inheritance of the RNAP-β gene in DST, Firmicutes and CPR. This correspondence further suggests that type 1 in DST, Firmicutes and CPR evolved by reduction through loss of b/lineage-specific BEAN insertions.

**Figure 6.**
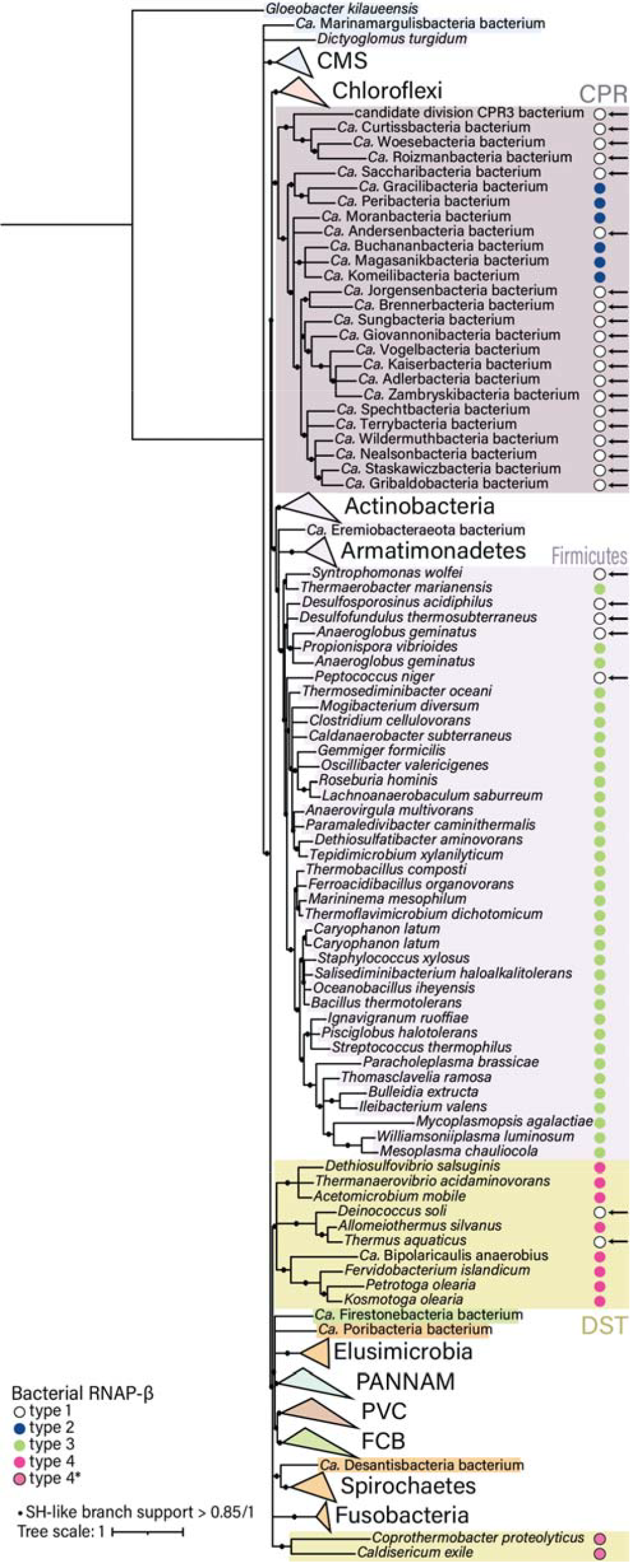
Maximum likelihood tree of RNAP-β. The ML tree (inferred with model Q.yeast +G+I) of RNAP-β was calculated using positions conserved in all bacteria.

Type 4 insertions in Terrabacteria may be convergent with type 4 insertions in Gracilicutes. Two scenarios could explain the distributions of type 4 RNAP-β insertions in two Terrabacteria groups, Armatimonadetes and DST. Type 4 RNAP-β insertions could arise from HGT from Gracilicutes or from vertical inheritance with convergence of the insertion site. In the case of Armatimonadetes, the first scenario (HGT from Gracilicutes) is not supported by our gene tree of RNAP-β (Figure 6 and Figure S4). Our gene tree is consistent with bacteria phylogenies that have been calculated using distinct gene markers (Coleman, et al. 2021; Megrian, et al. 2022; Witwinowski, et al. 2022). Thus, RNAP-β appears to have been vertically inherited to Armatimonadates and the location of its BEAN insertions appears to be convergent with type 4 RNAP-β from Gracilicutes. The location of BEAN insertions in RNAP-β from the DST group could also be convergent. However, in our gene tree, the DST appear as a sister lineage of Gracilicutes. Because RNAP-β from DST does not group within any of the Gracilicutes lineages, but appears as a separate sister lineage, the topology of our tree does not allow us to rule out or support a HGT of type 4 RNAP-β.

## Discussion

The data presented here are consistent with a model in which RNAP was subject to a discrete episode of aggressive domain insertion, around or after the last bacterial common ancestor, followed by a precipitous decline in the frequency of insertion (Figure 7). RNAP is a multi-subunit protein complex that contains RNAP-β and RNAP-β’ subunits. RNAP-β and RNAP-β’ are found in RNAPs in archaea, bacteria, eukarya, and nucleocytoplasmic large DNA viruses (Iyer, et al. 2001). Here we report that RNAP-β and RNAP-β’ each contain homologous insertional domains with idiosyncratic positions that generate block structures of RNAP-β and - β’ MSAs.

**Figure 7.**
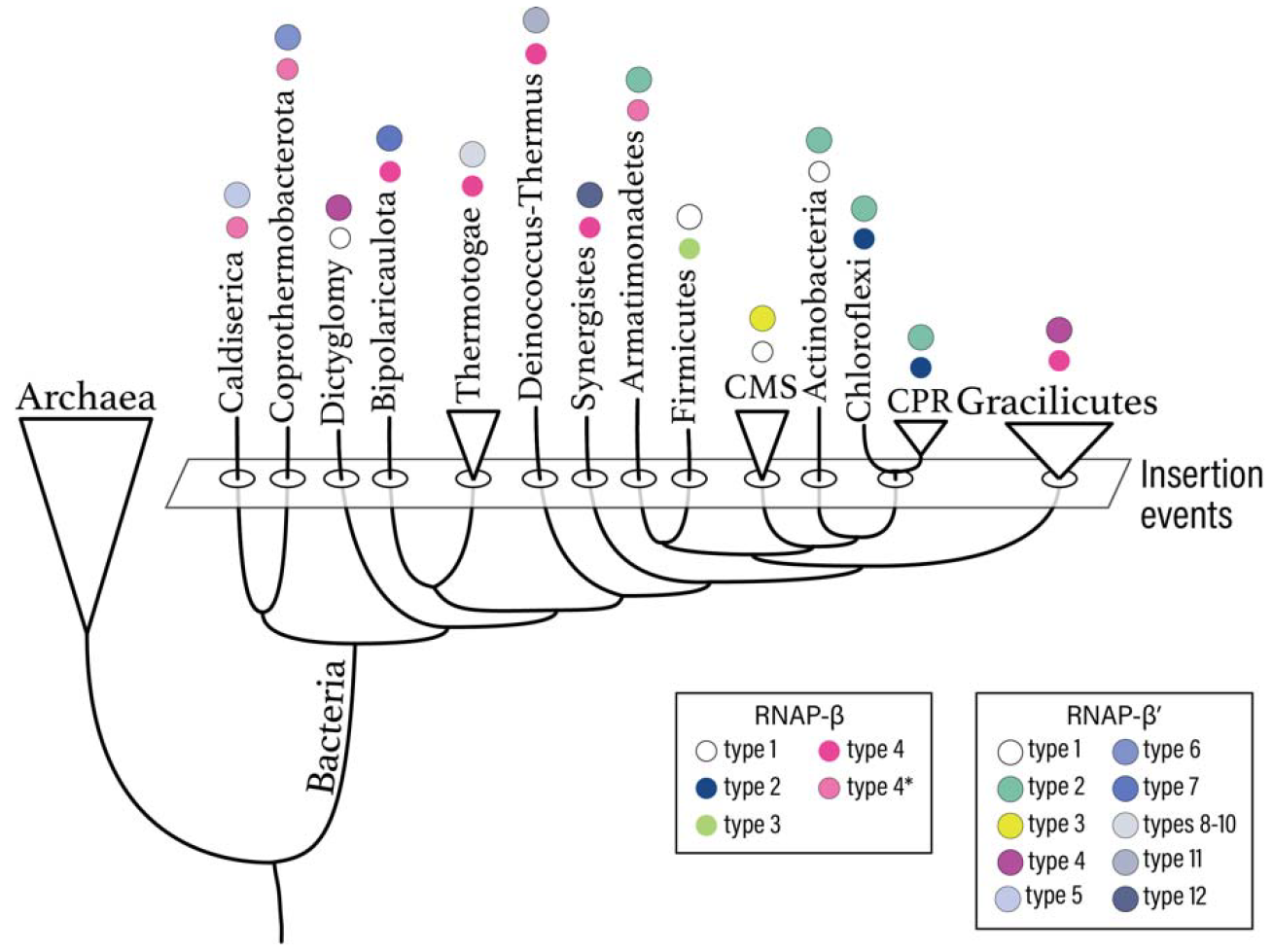
RNAP-β and RNAP-β’ domain insertions mapped into a schematic representation of the tree of bacteria. The tree reproduces the topology from Figures 5a and b, branches lengths have been altered. External nodes correspond to 14 bacterial lineages with 13 distinct combinations of RNAP-β and RNAP-β’ types.

Block structures of MSAs are not exclusive to RNAP-β and RNAP-β’ and have been described for universal components of the translation system [ribosomal proteins (Vishwanath, et al. 2004), and aminoacyl tRNA synthetases (Alvarez-Carreño, et al. 2023)]. But block differences in the translational system are observed between archaea and bacteria whereas here, in RNAP, they are observed within archaeal and bacterial domains. These insertional domains are informative about evolutionary history and pose important questions about evolutionary mechanisms.

We call the most common insertional domains BEAN and HABAS. The BEAN domain has a characteristic three-dimensional structure composed of two square bracket-like elements that are antiparallel relative to each other. Each bracket-like element is formed by an α-helix and two β-strands. The orientation between consecutive secondary elements is 90° within each bracket. The HABAS domain contains a four-stranded open β-sheet with a GD-box-like motif in one of the β-strands and the adjoining loop. In some instances, recursively inserted HABAS domains form complex domain-swapped structures.

### b/lineage-specific HABAS and BEAN domain insertions are polyphyletic

Insertional domains interrupt universal domains of RNAP-β and RNAP-β’, and thus, post-date the establishment of the basic multi-domain architectures of RNAP-β and RNAP-β’. We clustered RNAP-β and RNAP-β’ within bacteria based on number and kind of insertional domains. Insertions occur in distinct locations, allowing us to establish ‘types’ of RNAP-β and RNAP-β’. BEAN insertions specify the type of RNAP-β. HABAS domains specify the type of RNAP-β’.

Acquisition of BEAN domains may have been independent of acquisition of HABAS domains. We observe a mismatch in the distributions of RNAP-β and RNAP-β’ (Figures 5 and 7). For instance, bacteria from the DST group with type 4 RNAP-β have RNAP-β’ insertion sites that fall outside of our classification, but Armatimonadetes with type 4* RNAP-β have type 2 RNAP-β’, and Gracilicutes with type 4 RNAP-β have type 4 RNAP-β’. Below we discuss the phylogenetic distributions of RNAP-β and RNAP-β’ separately.

#### BEAN insertions and RNAP-β evolution

The distribution of RNAP-β types suggests that BEAN domains were inserted in the ancestors of three early branching bacterial lineages (Figure 7). These lineages are: i) the ancestor of Firmicutes, which acquired type 2 insertions; ii) the ancestor of Chloroflexi and CPR bacteria, which acquired type 3 insertions; and iii) the ancestor of all Gracilicutes, which acquired type 4 insertions. Armatimonadetes and DST also have type 4 insertions. Type 4 RNAP-β insertions could have arisen from HGT from Gracilicutes or from vertical inheritance with convergence of the insertion site. Convergence in BEAN locations suggest common characteristics of genome context that could have influenced the insertion sites.

#### HABAS insertions and RNAP-β’evolution

HABAS insertions in bacterial RNAP-β’ appear to have occurred in three ancestral populations: i) the ancestor of Armatimonadetes, Actinobacteria, Chloroflexi and CPR bacteria; ii) an ancestor within the CMS group; and iii) the ancestor of Gracilicutes. Extensive insertional diversity with the DST group suggests that these insertions occurred very early in bacterial evolution. The lack of insertional diversity in RNAP in late divergent groups suggests cessation of insertions later in bacterial evolution. The genes for RNAP-β and RNAP-β’ recorded and preserved the marks of evolutionary events that affected ancestral groups.

### Insertional domains in the evolution of translation and transcription

Our sequence similarity searches indicate that HABAS and BEAN are inserted in multiple unrelated proteins. We identify a b-specific BEAN insertion in RNAP-β’ and b/lineage specific BEAN insertions in bacterial RNAP-β. We observe BEAN insertions in the archaeal version of uL10. We identify a universal HABAS and a b-specific HABAS in RNAP-β and b/lineage specific HABAS insertions in RNAP-β’. Finally, we observe HABAS insertions in NusG, the only universally conserved transcription elongation factor (Werner and Grohmann 2011). HABAS insertions in NusG were identified in only bacteria from the DST group: *Fervidobacterium islandicum, Petrotoga olearia, Kosmotoga olearia* and *Candidatus* Bipolaricaulis anaerobius (Tables S1). The observation of a BEAN domain in the archaeal but not in the bacterial version of universal ribosomal protein L10 suggests some frequency of insertion before the last universal common ancestor (LUCA).

The local gene neighbourhood may have influenced the acquisition of BEAN and HABAS domains. The genes for RNAP-β and -β’ are adjacent to each other in the genomes of virtually all bacteria and most are in the neighbourhood of the genes that encode for NusG and universal ribosomal proteins uL1 and uL11 (Tables S1-S2). Similarly, in most archaea, uL10 is in the neighbourhood of the genes that encode for NusG, uL1 and uL11 (Table S3).

Transcription and translation are two of the central biological processes responsible for the encoding and synthesis of proteins. The patterns of insertion of HABAS and BEAN domains in universal and ancient proteins pose provocative questions regarding the timing and order of events during the early evolution of life. The mechanism of insertion remains unclear.

The combined data suggest that the bulk of the acquisition of BEAN domains in RNAP-β and archaeal uL10 and HABAS domains in RNAP-β’ occurred in ancestral lineages, shortly after LUCA, and that the descendants generally retained these insertions. We speculate that BEAN and HABAS insertions could be influenced by the genomic context. The slight differences in the locations of BEAN and HABAS insertions in RNAP-β and RNAP-β’ may reflect distinct bacterial lineages with distinct gene locations. Thus, in our model, b/lineage specific occurred in the deep evolutionary past, and just after an early divergence of the Last Bacterial Common Ancestor into distinct bacterial groups. The patterns that we observe left a mark on some of the first ancestral bacterial groups, and hint to an early diversification of Terrabacteria, particularly of the DST group.

## Methods

### Identification of RNAP-β and β’ subunits in bacteria and archaea

The sequences of RNAP subunits β and β’ from *Sulfolobus acidocaldarius* (UniProt IDs: P11513 and P11512) and *Bacillus subtitlis* (UniProt IDs: P37870 and P37871) were searched in a set of archaeal and bacterial proteomes derived from (Moody, et al. 2022) using phmmer from the HMMER3 suite (Eddy 2011). Sequences above threshold (E-value < 1×10^-10^) were retrieved and aligned. Multiple sequence alignments (MSAs) were generated with the einsi option from MAFFT v7 (Katoh and Standley 2013).

### Domain annotation of RNAP-β and β’ and classification

The MSAs of RNAP subunits β and β’ were converted each into a sequence profile and compared to CATH_S40, ECOD_F70 and SCOPe95 with HH-search (Steinegger, Meier, et al. 2019) on the MPI Bioinformatics Toolkit (Zimmermann, et al. 2018). CATH_S40 contains CATH domains clustered at 40% sequence identity; ECOD_F70 contains ECOD domains clustered at 70% sequence identity; and SCOPe95 contains domain sequences clustered at 95% sequence identity. The block patterns on the MSAs were used as reference to classify the multi-domain organization types of bacterial RNAP-β and RNAP-β’ proteins (Figure S1). For each RNAP-β and RNAP-β’ type, a representative was selected for structure analysis. All representatives have a experimentally determined structure in the PDB (Berman, et al. 2000) or a predicted structure in AlphaFold DB (Varadi, et al. 2022). Per-residue confidence score (pLDDT) and predicted aligned error (PAE) of the structure predictions are shown in Figures S6-S16. Sequences with unique insertion patterns were annotated individually, and the annotations were inspected over structure predictions generated with AlphaFold version 2.0 (Jumper, et al. 2021).

### Identification of HABAS and BEAN domains homologs

A-, b- and b/lineage-specific insertions were trimmed according to the blocks in the MSAs. Profiles were calculated for each trimmed MSA with hhmake from the hh-suite (Steinegger, Meier, et al. 2019), considering columns with fewer than 50% gaps match states.

The MSAs were converted to profile Hidden Markov Models using the HH-suite version 3.3.0 (Steinegger, Meier, et al. 2019). The profiles were searched against the BFD database (Steinegger, Mirdita, et al. 2019) of clustered genome and metagenome sequences using HHblits (3 iterations, Probability > 60). Significant matches (minimum probability: 60, minimum coverage with master sequence 80%) were retrieved and clustered with CLANS (Gabler, et al. 2020) by all-against-all BLASTP sequence similarity (P-value 1×10-20). Groups with at least 30 homologs in the cluster map were extracted; realigned with MAFFT einsi (Katoh and Standley 2013); and converted to HMM profiles with HMMER version 3.3.2 (Eddy 2011). The HMM profiles were searched with phmmer in the same set of archaeal and bacterial proteomes (Moody, et al. 2022) that was used RNAP-β and RNAP-β’ identification.

Structure based MSAs of HABAS and BEAN domains were calculated with MATRAS (Kawabata 2003).

### Maximum likelihood tree of RNAP-β

The MSA of bacteria RNAP-β was trimmed with trimAl v1.4 to remove positions with more than 10% gaps. The ML tree was calculated with PhyML (Guindon, et al. 2010) on the Montpellier Bioinformatics Platform. Model selection was determined with SMS (Lefort, et al. 2017).

## Data availability

Sequence alignments and structure predictions associated with this manuscript have been deposited in the FigShare repository DOI: 10.6084/m9.figshare.25663923.

## Acknowledgements

The authors thank Finn Werner for insightful discussions. This work was funded by the National Aeronautics and Space Administration grant 80NSSC18K1139. Claudia Alvarez-Carreño’s research was supported by the Royal Society Newton International Fellowship.

