## Supplementary material for "BEAN and HABAS: Polyphyletic insertions in RNAP that point to deep-time evolutionary divergence of bacteria": Figures S1-S16

* Corresponding authors:

Claudia Alvarez-Carreño

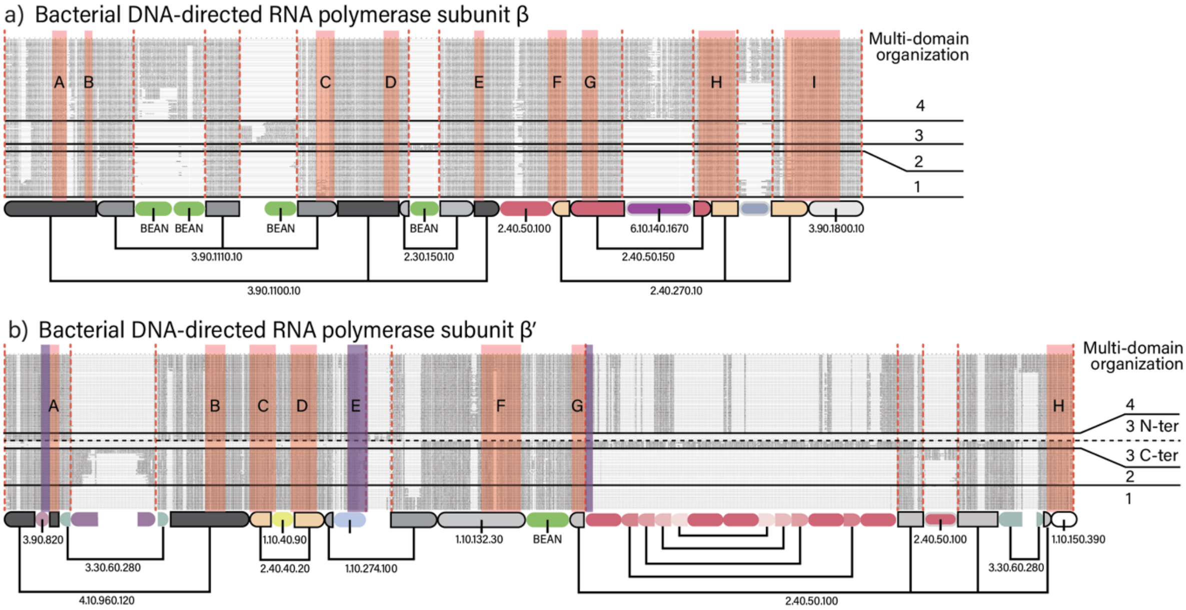


**Supplementary Figure 1. Blocks of conservation in the multiple sequence alignments of bacterial RNAP-β and RNAP-β’.** a) Multiple sequence alignment of RNAP-β and domain annotation. b) Multiple sequence alignment of RNAP-β and domain annotation. Sequences are grouped by type. Vertical lines delineate the blocks of differential conservation: universal; archaeal; bacterial; and bacterial lineage specific. Horizontal lines delineate the different types. Conserved sequence motifs as defined by (Sweetser, et al. 1987; Jokerst, et al. 1989) are indicated by shaded bands. Red: universally conserved sequence motifs. Purple: bacteria-specific sequence motifs.

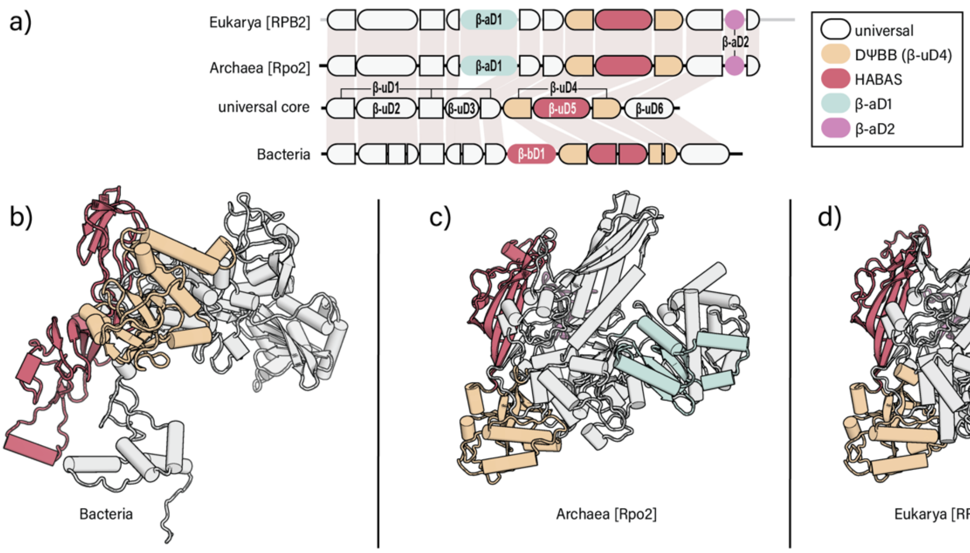


**Supplementary Figure 2. Multi-domain organizations of RNAP-β in bacteria, archaea and eukarya.** (a) Domain organization of RNAP-β. First row: domains of eukaryotic RNAP-β. Second row: domains of archaeal RNAP-β. Third row: universally shared domains between archaeal and bacterial orthologs. Fourth row: domains of bacterial type 1 RNAP-β. (b to e) Three-dimensional structure of the RNAP-β orthologs. Domains are coloured as in (a).

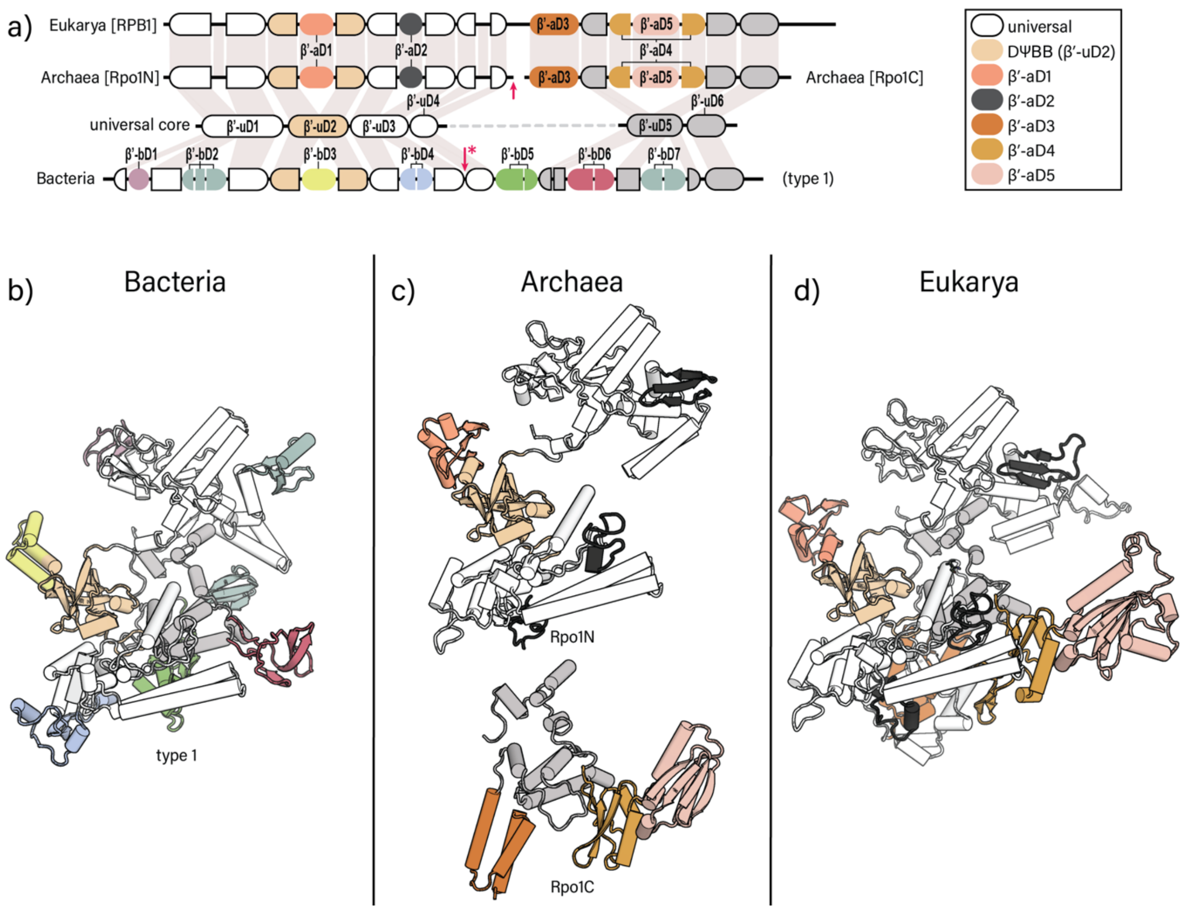


**Supplementary Figure 3. Multi-domain organizations of RNAP-β' in bacteria, archaea and eukarya.** (a) Domain organization of RNAP-β'. First row: domains of eukaryotic RNAP-β'. Second row: domains of archaeal RNAP-β'. Third row: universally shared domains between archaeal and bacterial orthologs. Fourth row: domains of bacterial type 1 RNAP-β'. (b to e) Three-dimensional structure of the RNAP-β' orthologs. Domains are coloured as in (a).

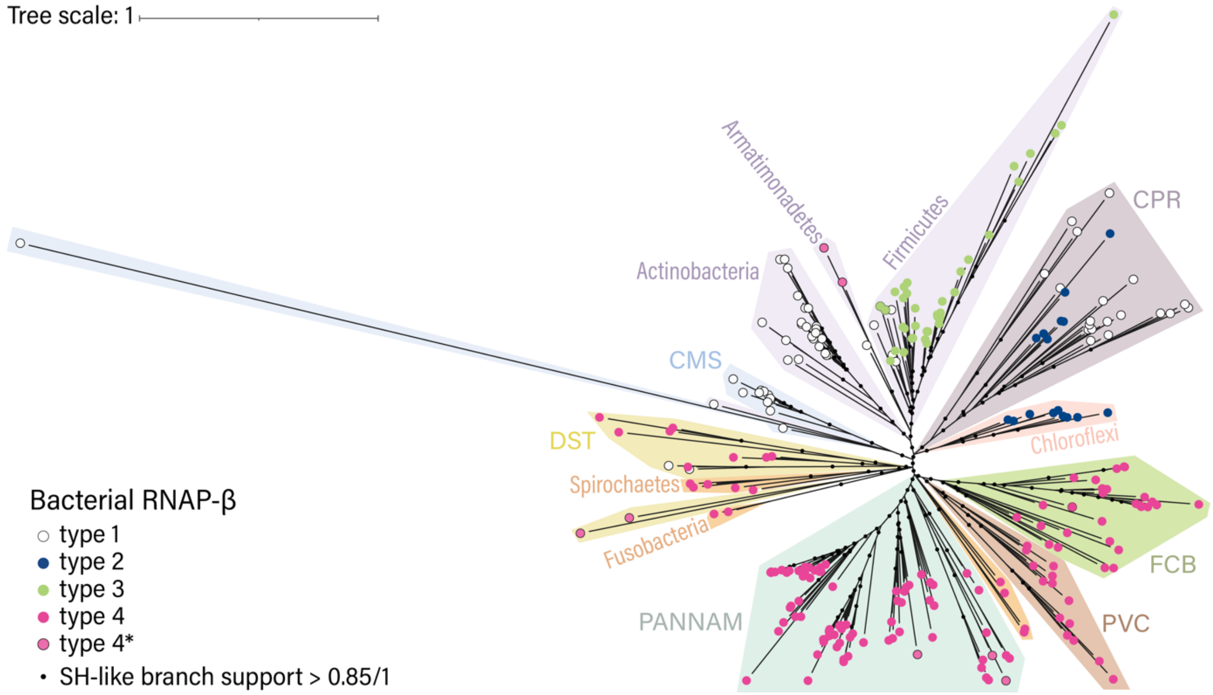


**Supplementary Figure 4. Maximum likelihood tree of RNAP-β.** The ML tree (inferred with model Q.yeast +G+I) of RNAP-β was calculated using positions conserved in at least 90% of sequences in the MSA.

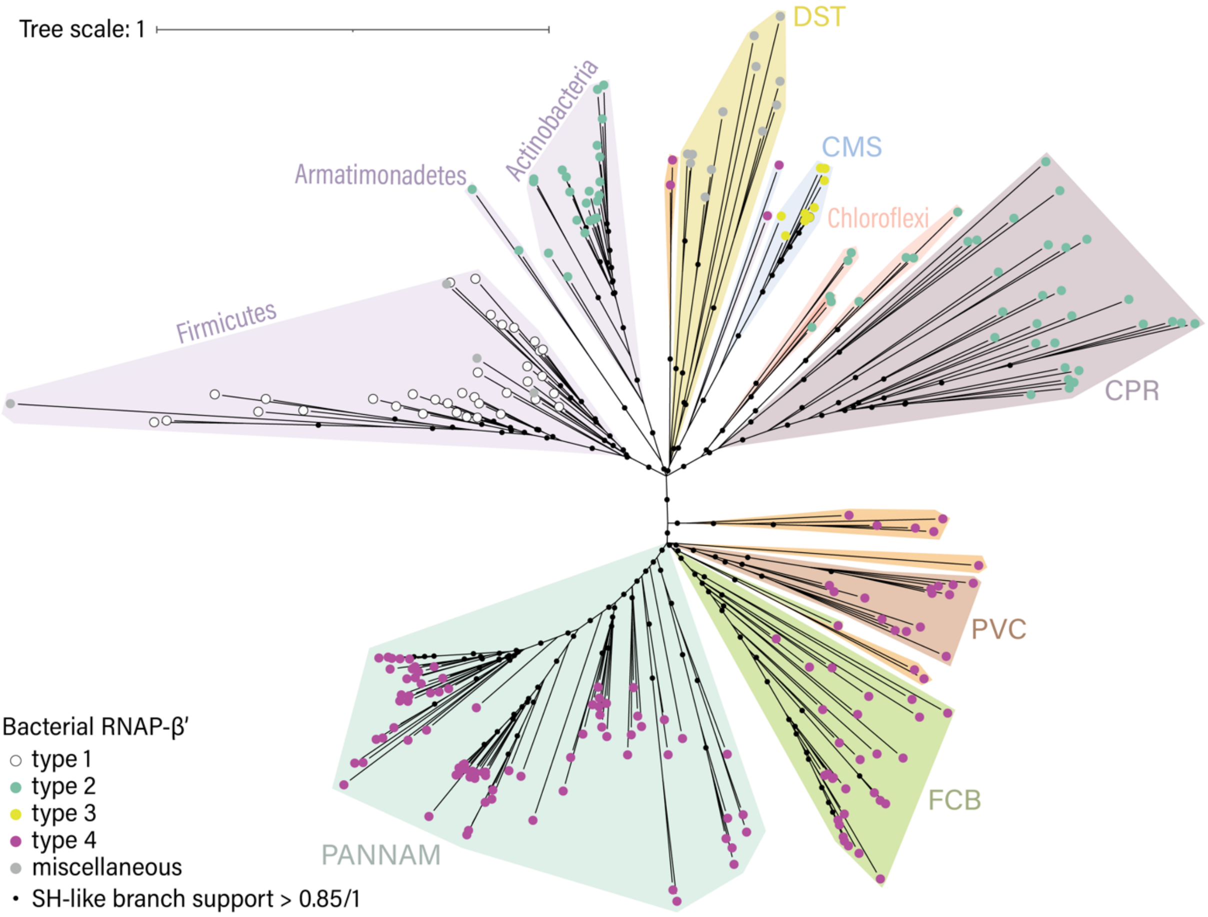


**Supplementary Figure 5. Maximum likelihood tree of RNAP-β'.** The ML tree (inferred with model LG+G+I) of RNAP-β' was calculated using positions conserved in at least 90% of sequences in the MSA. Type 3 RNAP-β'$_{BacN}$ and RNAP-β'$_{BacC}$ sequences are concatenated in the MSA.

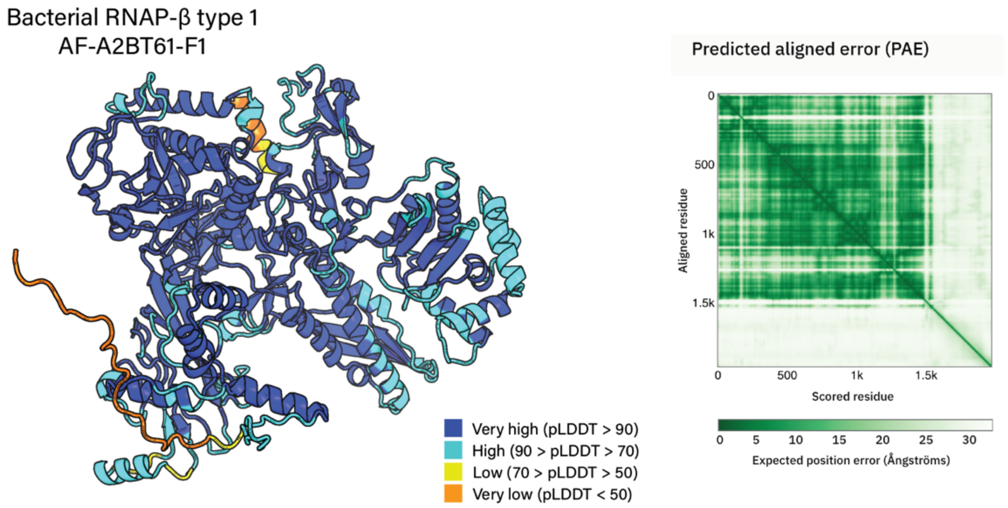


**Supplementary Figure 6. Structure prediction of type 1 RNAP-β from *Prochlorococcus marinus***. The structure prediction was obtained from AlphaFold DB (Varadi, et al. 2022)(ID: AF-A2BT61-F1).

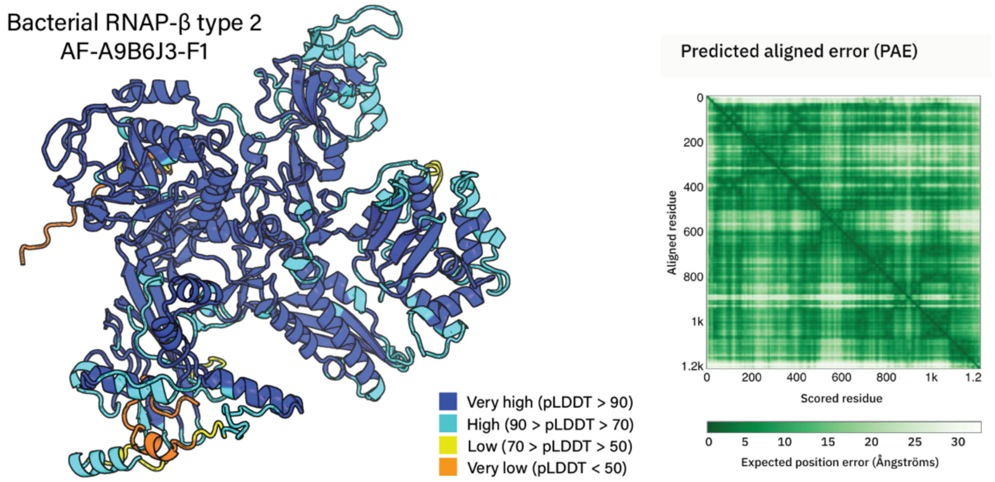


**Supplementary Figure 7. Structure prediction of type 2 RNAP-β from *Herpetosiphon aurantiacus***. The structure prediction was obtained from AlphaFold DB (Varadi, et al. 2022)(ID: AF-A9B6J3-F1).

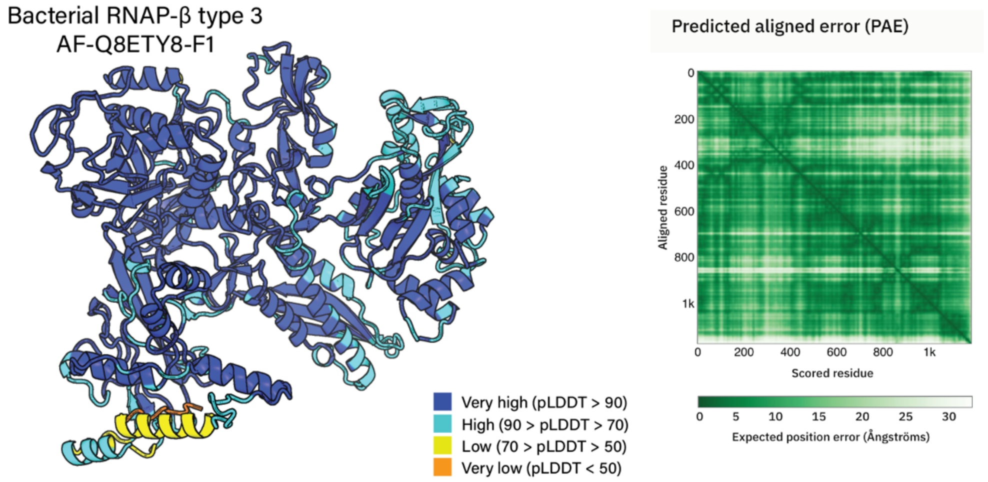


**Supplementary Figure 8. Structure prediction of type 3 RNAP-β from *Oceanobacillus iheyensis***. The structure prediction was obtained from AlphaFold DB (Varadi, et al. 2022)(ID: AF-Q8ETY8-F1).

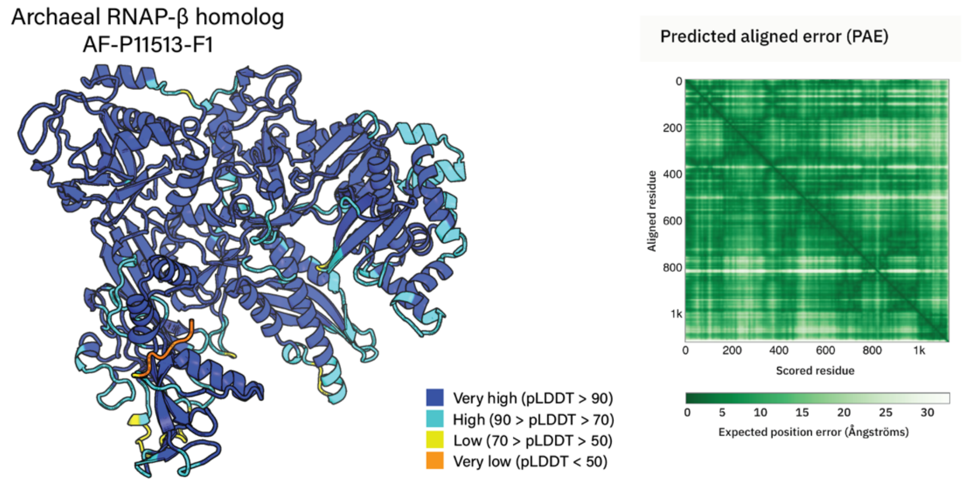


**Supplementary Figure 9. Structure prediction of the archaeal homolog of RNAP-β from *Sulfolobus acidocaldarius*.** The structure prediction was obtained from AlphaFold DB (Varadi, et al. 2022)(ID: AF-P11513-F1).

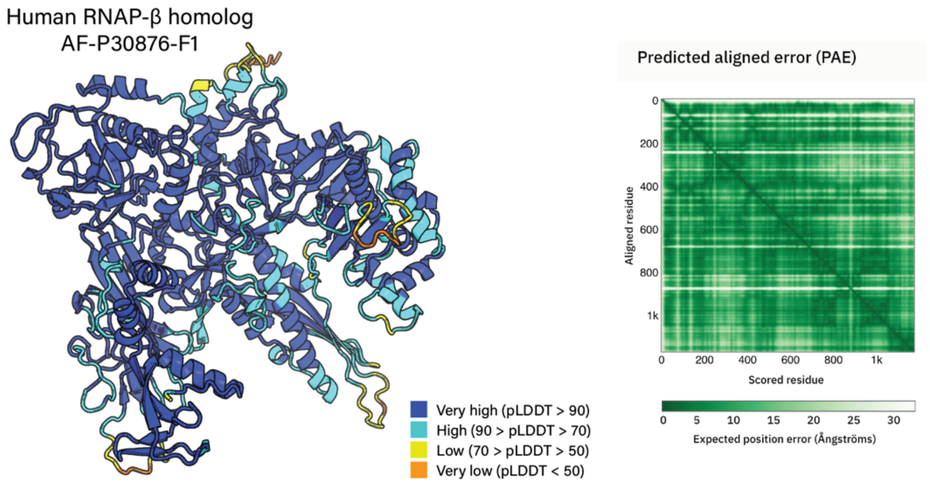


**Supplementary Figure 10. Structure prediction of the eukaryotic RNAP-β homolog from *Homo sapiens***. The structure prediction was obtained from AlphaFold DB (Varadi, et al. 2022)(ID: AF-P30876-F1).

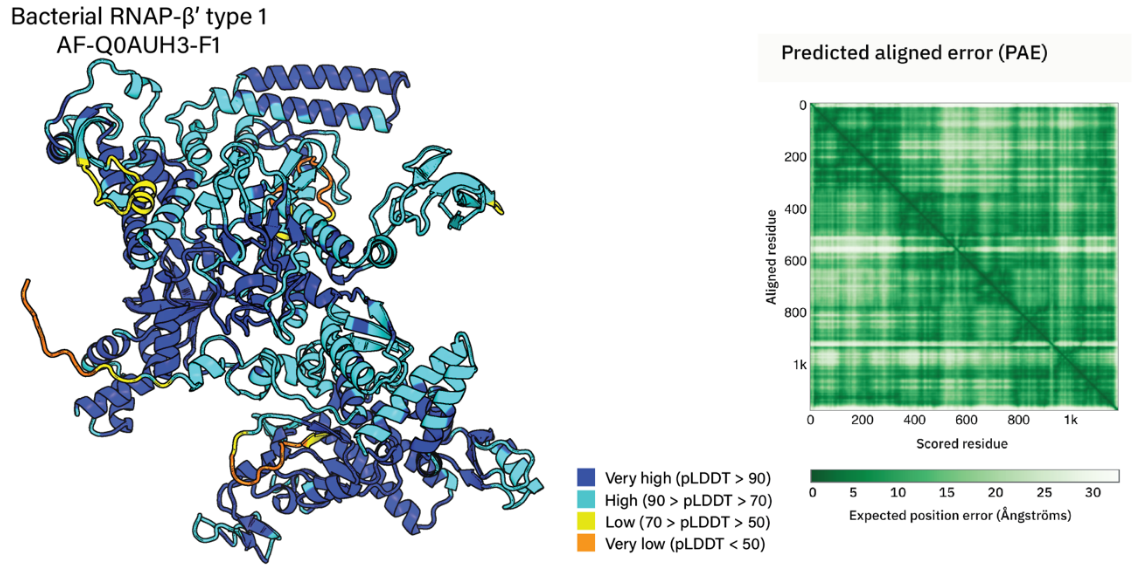


**Supplementary Figure 11. Structure prediction of type 1 RNAP-β' from *Syntrophomonas wolfei* subsp. *wolfei***. The structure prediction was obtained from AlphaFold DB (Varadi, et al. 2022)(ID: AF-Q0AUH3-F1).

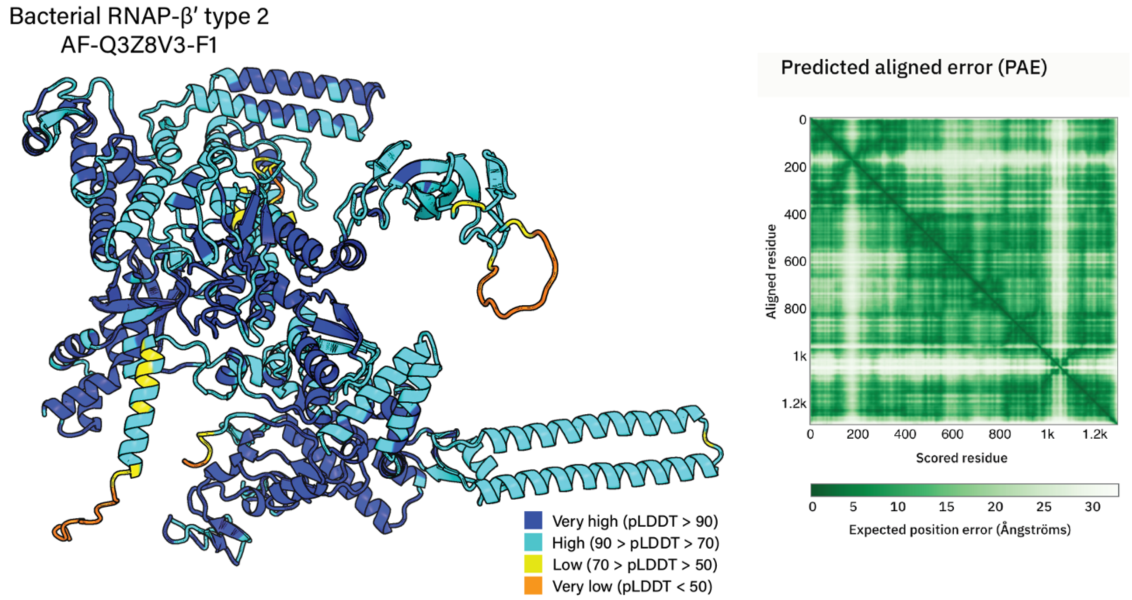


**Supplementary Figure 12. Structure prediction of type 2 RNAP-β' from *Dehalococcoides mccartyi***. The structure prediction was obtained from AlphaFold DB (Varadi, et al. 2022)(ID: AF-Q3Z8V3-F1).

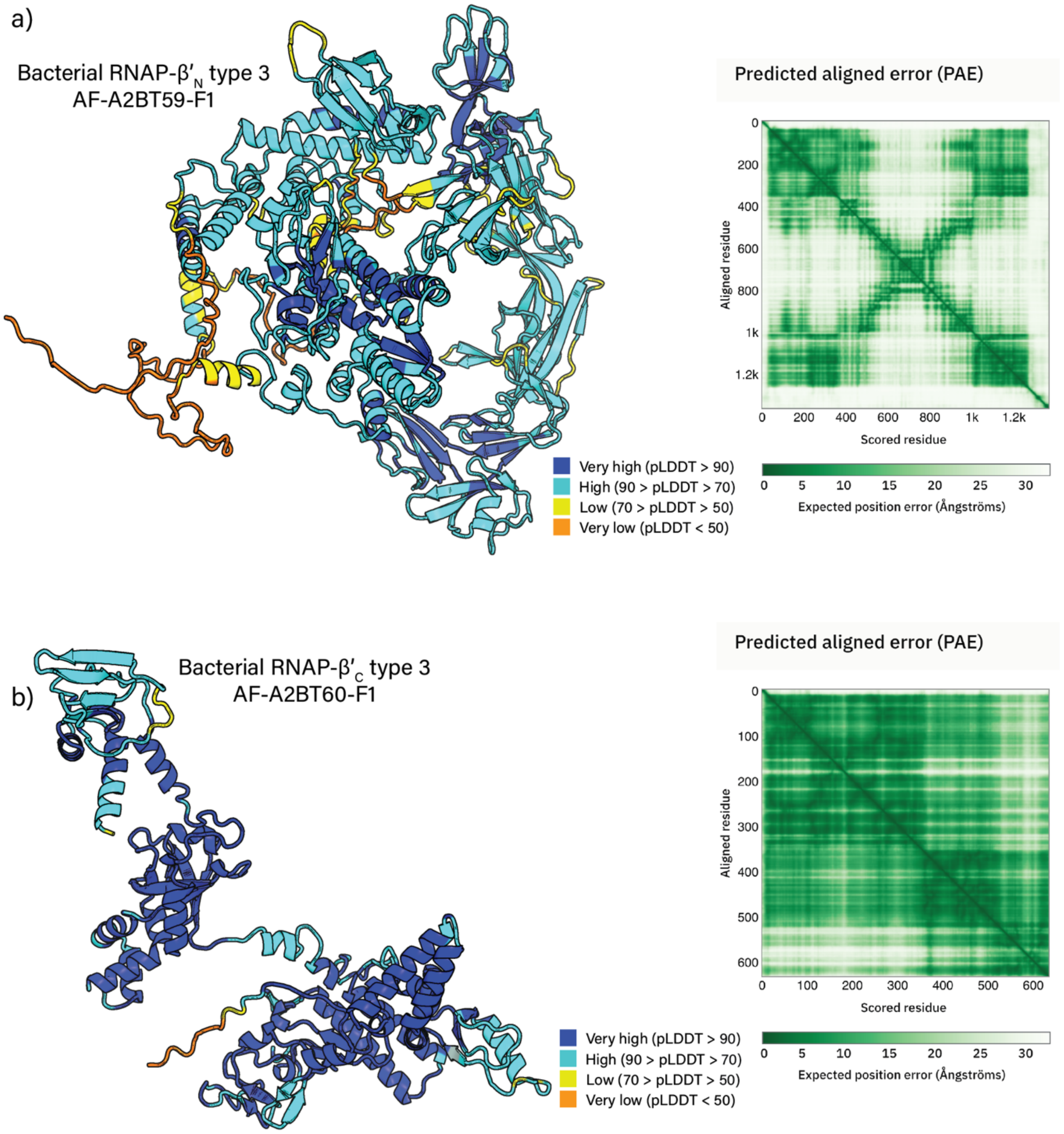


**Supplementary Figure 13. Structure prediction of type 3 RNAP-β' from *Prochlorococcus marinus***. The structure predictions were obtained from AlphaFold DB (Varadi, et al. 2022) (a) RNAP-$\beta'_N$ (AF-A2BT59-F1). (b) RNAP-$\beta'_C$ (AF-A2BT60-F1).

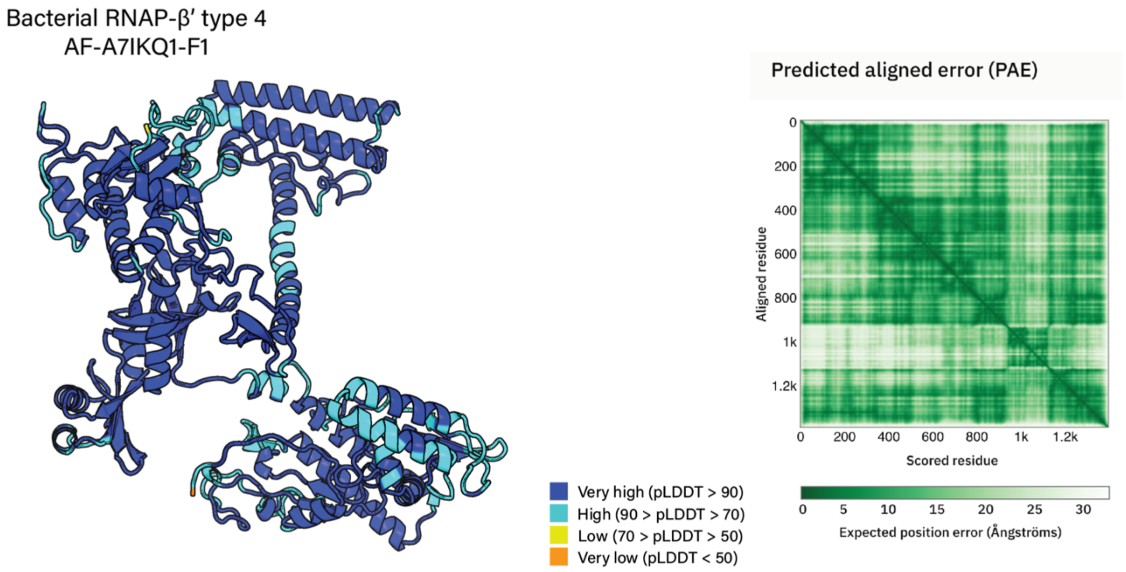


**Supplementary Figure 14. Structure prediction of type 4 RNAP-β' from *Xanthobacter autotrophicus***. The structure prediction was obtained from AlphaFold DB (Varadi, et al. 2022)(ID: AF-A7IKQ1-F1).

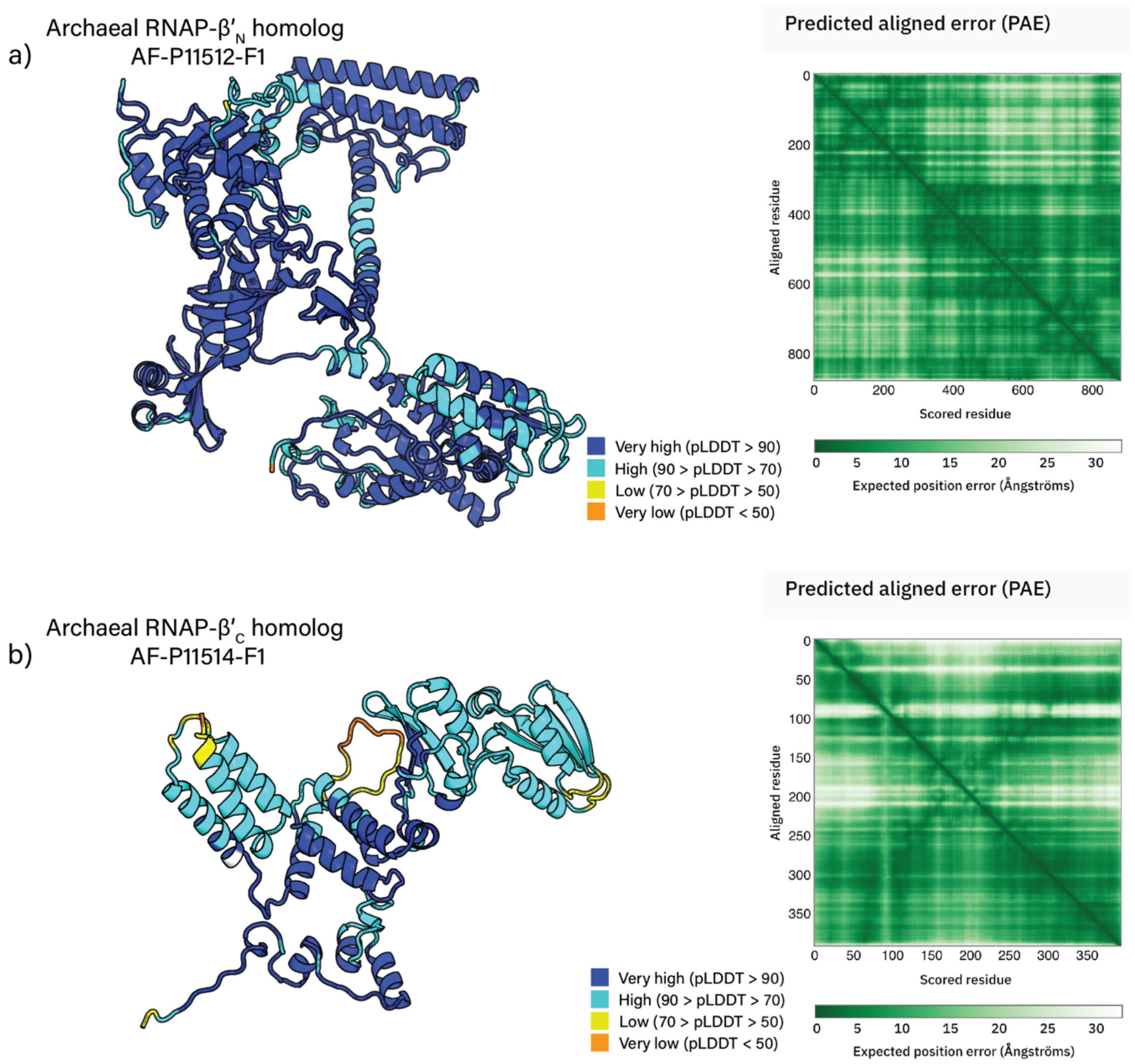


**Supplementary Figure 15. Structure prediction of the archaeal ortholog of RNAP-β' from *Sulfolobus acidocaldarius***. The structure predictions were obtained from AlphaFold DB (Varadi, et al. 2022) (a) RNAP-$\beta'_N$ (AF-P11512-F1). (b) RNAP-$\beta'_C$ (AF-P11514-F1).

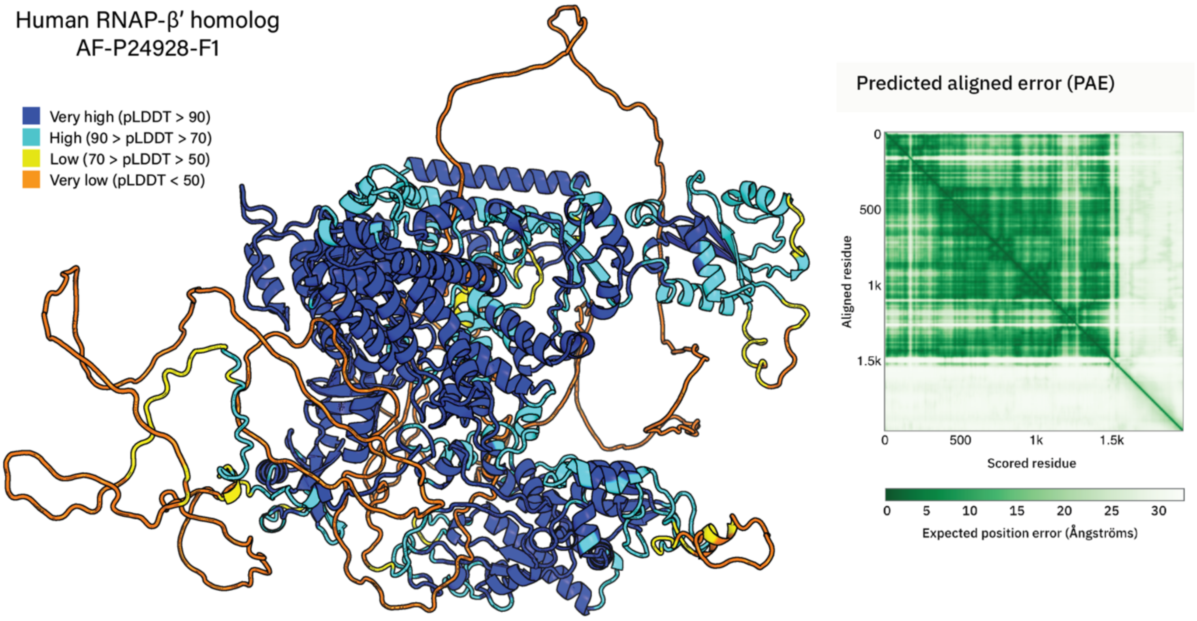


**Supplementary Figure 16. Structure prediction of an eukaryotic ortholog of RNAP-β′ from *Homo sapiens***. The structure prediction was obtained from AlphaFold DB (Varadi, et al. 2022)(ID: AF-P24928-F1).
